# Modulation of *Prdm9*-controlled meiotic chromosome asynapsis overrides hybrid sterility in mice

**DOI:** 10.1101/203505

**Authors:** Sona Gregorova, Vaclav Gergelits, Irena Chvatalova, Tanmoy Bhattacharyya, Barbora Valiskova, Vladana Fotopulosova, Petr Jansa, Diana Wiatrowska, Jiri Forejt

## Abstract

The infertility of hybrids between closely related species is one of the reproductive isolation mechanisms leading to speciation. *Prdm9*, the only known vertebrate hybrid sterility gene causes failure of meiotic chromosome synapsis and infertility in male hybrids between mouse strains derived from two mouse subspecies. Within species *Prdm9* determines the sites of programmed DNA double-strand breaks and meiotic recombination hotspots. To investigate the relation between *Prdm9*-controlled meiotic arrest and asynapsis, we inserted random stretches of consubspecific homology on several autosomal pairs in sterile hybrids and analyzed their ability to form synaptonemal complexes and rescue male fertility. Twenty-seven or more Mb of consubspecific homology fully restored synapsis in a given autosomal pair and we predicted that two symmetric DSBs or more per chromosome are necessary for successful meiosis. We hypothesize that impaired recombination between evolutionary diverged homologous chromosomes could function as one of the mechanisms of hybrid sterility occurring in various sexually reproducing species.

## Introduction

Hybrid sterility (HS) is a postzygotic reproductive isolation mechanism that safeguards speciation by restricting gene flow between related taxa. HS is a universal phenomenon observed in many eukaryotic inter-species hybrids, including yeast, plants, insects, birds, and mammals (Coyne and Orr 2004; Maheshwari and Barbash 2010). In the early days of genetics, HS had been difficult to accommodate in Darwin's theory of evolution by natural selection until the Bateson - Dobzhansky - Muller incompatibility (BDMI) hypothesis (Muller and Pontecorvo 1942; Dobzhansky 1951; Orr 1996) explicated HS and, more generally, any hybrid incompatibility, as a consequence of the independent divergence of mutually interacting genes and resulting aberrant interaction of the new alleles, previously not tested by natural selection. HS has several common features across various sexually reproducing eukaryotic species. Haldane's rule posits that if one sex of the F1 offspring of two different animal races is absent, rare, or sterile, it is the heterogametic sex (XY or ZW) (Haldane 1922). Another common feature refers to the disproportionately large role of Chr X compared to autosomes in reproductive isolation (Presgraves 2008). More recently, interaction between selfish genomic elements causing meiotic drive and their suppressors has been implicated in some instances of reproductive isolation (Orr 2005; Zhang *et al.* 2015)

The molecular mechanisms underlying HS remain an unresolved question. Historically, genic and chromosomal mechanisms of HS were hypothesized, but the latter were soon dismissed as unlikely on the grounds that large chromosomal rearrangements do not segregate with HS genetic factors (Dobzhansky 1951). Other possible forms of non-genic chromosomal HS were not considered because of the limited knowledge of the carrier of genetic information at the time. Thus, for the last 80 years or so, the focus on the genic control of HS prevailed (Dobzhansky 1951; Orr 1996; Forsdyke 2017). In studies mapping HS genes, the *Drosophila* group of species has been the model of choice, yet only five *Drosophila* HS genes, namely *OdsH, JYAlpha, Ovd, agt*, and *Taf1*, have been identified so far, none of which has a known interacting gene predicted by the BDMI hypothesis (Ting *et al.* 1998; Masly *et al.* 2006; Phadnis and Orr 2009). The low success rate of the positional cloning of HS genes was explained by the oligogenic or polygenic nature of HS phenotypes and by the inherent difficulty in genetically dissecting the phenotype that prevents its own transfer to progeny.

Over forty years ago, we introduced the house mouse (*Mus musculus*) as a mammalian model for the genetic analysis of HS. The first HS locus *Hst1* was genetically mapped in crosses of laboratory inbred strains (predominantly of *Mus musculus domesticus* (*Mmd*) origin) with wild *Mus musculus musculus* (*Mmm*) mice (Forejt and Ivanyi 1974). Later, we developed PWD/Ph and PWK/Ph inbred strains purely from the wild *Mmm* mice of Central Bohemia (Gregorova and Forejt 2000) and used them in the positional cloning of *Hst1* by high-resolution genetic crosses and physical mapping (Gregorova *et al.* 1996; Trachtulec *et al.* 1997). Finally, we identified the *Hst1* locus with the PR domain containing 9 gene (*Prdm9*) (Mihola *et al.* 2009), coding for histone H3 lysine 4/lysine 36 methyltransferase (Powers *et al.* 2016) the first and still the only HS gene known in vertebrates. Most of the tested laboratory inbred strains share either the *Prdm9^Dom2^* or *Prdm9^Dom3^* allele. (Parvanov *et al.* 2010; Brunschwig *et al.* 2012). The former allele was found in inbred strains producing sterile male hybrids when crossed with PWD females, while the *Prdm9^Dom3^* was observed in the strains that yielded quasi-fertile males in the same type of intersubspecific crosses (Forejt *et al.* 2012).

The male sterility of (PWD x C57BL/6)F1 (henceforth PB6F1) hybrids depends on the interaction of the heterozygous allelic combination *Prdm9^Msc^*/*Prdm9^Dom2^* with the PWD allelic form of the X-linked Hybrid sterility X chromosome 2, *Hstx2^Msc^* locus (Dzur-Gejdosova *et al.* 2012; Bhattacharyya *et al.* 2014). For the sake of clarity and to stress the origin of the alleles we will use *Prdm9^PWD^*, *Prdm9^B6^* and *Hstx2^PWD^* in the rest of this paper. Any other tested allelic combination of these two major HS genes yields fully fertile or subfertile male hybrids (Dzur-Gejdosova *et al.* 2012; Flachs *et al.* 2012). The proper allelic combination of *Prdm9* and *Hstx2* genes is necessary but not sufficient to completely govern HS because less than 10% instead of expected 25% of (PWD x B6) x B6 male backcross progeny replicated the infertility of male PB6F1 hybrids (Dzur-Gejdosova *et al.* 2012). Initially, we explained this ‘missing heritability’ by assuming additional genic interaction of three or more additional HS genes with a small effect that escaped the genetic screen (Dzur-Gejdosova *et al.* 2012). However, an alternative, non-genic explanation emerged from the analysis of meiotic phenotypes of sterile hybrids. The failure of multiple autosomal pairs to synapse properly, the persistence of phosphorylated histone γH2AX indicating unrepaired DNA double-strand breaks (DSBs) on unsynapsed chromosomes, and the disturbed transcriptional inactivation of sex chromosomes at the first meiotic prophase were the most prominent features observed in approximately 90% of primary spermatocytes of infertile PB6F1 inter-subspecific hybrids (Bhattacharyya *et al.* 2013; Bhattacharyya *et al.* 2014). We suggested that the failure of chromosomes to synapse could be related to their fast-evolving nongenic DNA divergence. Because failure of proper synapsis in pachytene spermatocytes is known to interfere with the normal progression of the first meiotic division (Mahadevaiah *et al.* 2008; Burgoyne *et al.* 2009), we assumed that the asynapsis could be the ultimate cause of the sterility of male hybrids. Recently, the role of PRDM9 zinc finger domain binding sites within noncoding genomic DNA has been demonstrated in PB6F1 male HS. Replacement of the mouse sequence encoding the PRDM9 zinc-finger array with the orthologous human sequence reversed sterility in (PWD × B6-*Prdm9^Hu^*)F1 hybrid males (Davies *et al.* 2016). In PB6F1 hybrids, roughly 70% of *Prdm9*-directed DSBs hotspots identified by the DMC1 ChIP-seq method were enriched on the ‘nonself’ homologous chromosome, since the DSBs determined by the B6 allele of *Prdm9* were found predominantly or exclusively on PWD chromosomes, and *vice versa*. Such hotspots were designated as asymmetric DSB hospots. Chromosome-specific quantification of asymmetry correlated well with the asynapsis rate across five arbitrarily chosen chromosomes of PB6F1 hybrids (Davies *et al.* 2016; Smagulova *et al.* 2016). Another, non-exclusive interpretation of DMC1 ChIP-seq data pointed to significant enrichment of PRDM9-independent hotspots in the PB6F1 hybrid testis, which occurs in promoters and other regulatory motifs and which is characteristic of spermatogenic arrest in *Prdm9* knockout males (Smagulova *et al.* 2016). Recently, one third of PRDM9-dependent DSBs was reported within sequences that have at least some repetitive character and indicating that inappropriately high DSB levels in retroposons and other repetitive elements may contribute to the infertility seen in some mouse hybrids (Yamada *et al.* 2017).

In this work, we studied the relationship between meiotic chromosome asynapsis, intersubspecific heterozygosity and male HS in a series of PB6F1 hybrids carrying recombinant chromosomes with *Mmm/Mmm* consubpecific (belonging to the same subspecies) PWD/PWD homozygous intervals on *Mmm/Mmd* intersubpecific (belonging to different subspecies) PWD/B6 heterozygous background. We report the restoration of synapsis of intersubspecific chromosome pairs in the presence of 27 Mb or more of consubspecific sequence, and the reversal of HS by targeted suppression of asynapsis in the four most asynapsis-sensitive chromosomes. Thus the asymmetry of PRDM9 hotspots is linked to failure of synapsis of homologous chromosomes and to meiotic arrest of sterile hybrid males.

## Results

### Small chromosomes are more susceptible to asynapsis in sterile F1 hybrids

First, we ascertained the frequency of meiotic asynapsis separately for each chromosome pair of PB6F1 hybrid males by combining fluorescence in-situ hybridization (FISH) to decorate chromatin from individual chromosomes with immunostaining of the synaptonemal complex protein 3 (SYCP3) a major component of axial/lateral elements to visualize synaptonemal complexes, and a HORMA domain-containing protein-2, HORMAD2 (Wojtasz *et al.* 2012), to identify the axial elements of unsynapsed chromosomes (*figure supplement 1A*). Altogether 4168 pachynemas from 40 PB6F1 hybrid males were analyzed. All autosomes of hybrid males displayed a certain degree of asynapsis, classified as complete, partial, or intermingled, with frequencies ranging from 2.6% (Chr 1) to 42.2% (Chr 19) (*Figure 1A - table supplement 1*). A strong bias was evident towards higher asynapsis rates in the five smallest autosomes (P < 5.3×10^−14^, comparison of GLMM models, *Figure 1A*). Recently, SPO11 oligos released during the processing of DSBs were sequenced, mapped and quantified at chromosome-wide scale in male mice of the B6 laboratory inbred strain (Lange *et al.* 2016). This information, together with the estimated frequency of asymmetric DSB hotspots in PBF1 hybrids (Davies *et al.* 2016; Smagulova *et al.* 2016) enabled us to calculate the possible correlation between the number of symmetric DSBs per chromosome and synapsis between intersubspecific homologs. The calculation is based on and limited by the following premises: (i) the overall densities of DSBs on individual chromosomes of B6 and PB6F1 hybrid males are similar, (ii) approximately 250 DSBs occur per leptotene/zygotene cell (Kauppi *et al.* 2013), and (iii) the 0.28 proportion of symmetric DSB hotspots in (PWD x B6)F1 hybrid males (Davies *et al.* 2016) is constant in all autosomes. Under these conditions, a strong negative correlation (Spearman’s ρ=-0.76, P = 0.0002) of asynapsis rate with symmetric DSB hotspots (Lange *et al.* 2016) can be seen (*Figure 1B - table supplement 2*). This correlation is stronger than the correlation of the asynapsis rate with the chromosomal physical length (Spearman’s ρ=-0.68, P = 0.0013). Importantly, the asynapsis rate cannot be better explained by the chromosomal length (P = 0.709, comparison of GLMM models), when it is controlled for the symmetric DSB hotspots (SPO11 oligos), while it is better explained by the symmetric DSB hotspots (SPO11 oligos) when controlled for chromosomal length (P = 0.046, comparison of GLMM models). Thus our findings indicate that synapsis of a pair of homologous chromosomes depends on the presence of a certain minimum number of symmetric DSB hotspots.

**Figure 1.**
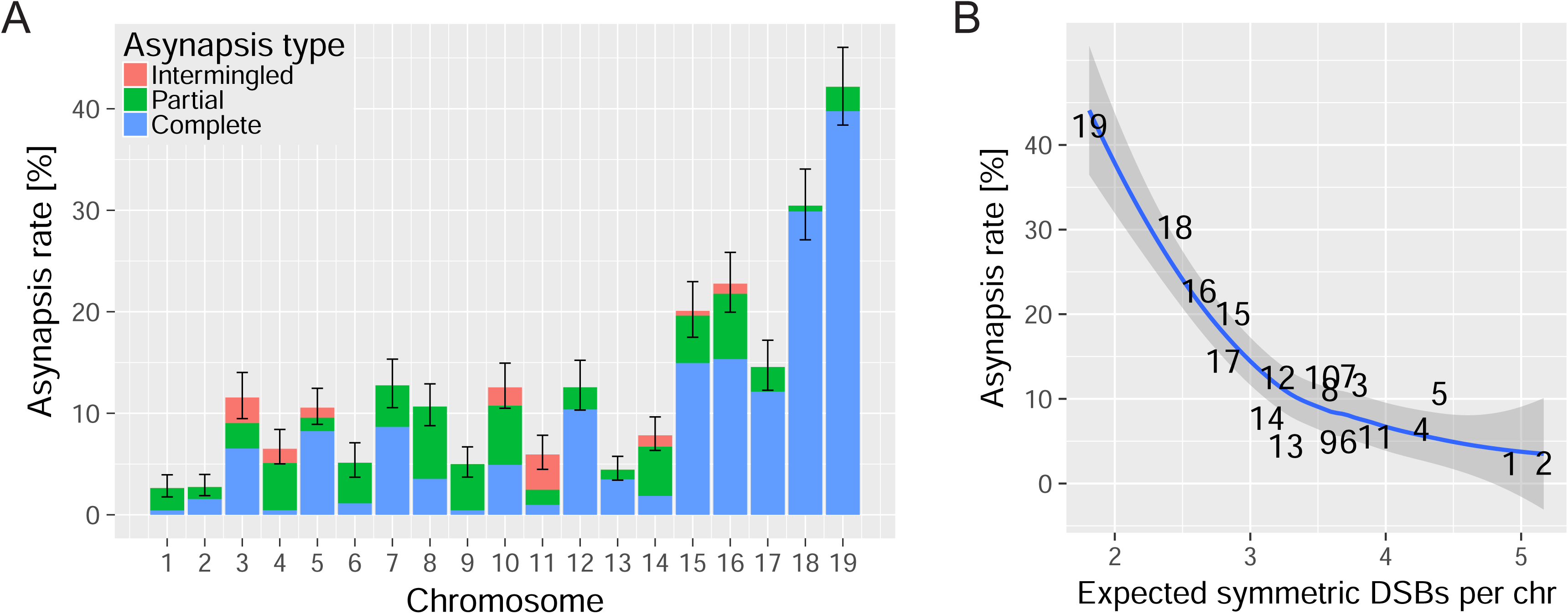
The asynapsis rate of individual autosomes in sterile male (PWD x B6)F1 hybrids. (*A*) Mean asynapsis rate ± S.E (based on GLMM model). Intermingled asynapsis refers to overlaps of two or more asynapsed chromosomes within the DNA FISH cloud of chromatin. Smaller chromosomes had higher asynapsis rate (GLMM model, P < 1.1×10^−13^). Concurrently, the chromosomes with higher asynapsis rate were also more involved in complete rather than partial asynapsis (GLMM model, P < 6.3×10^−5^). Proportion of complete and partial asynapsis was controlled by the asynapsis rate rather the chromosomal length (test for effect of the length when controlled for the asynapsis rate, 0.491). (*B*) Negative correlation (Spearman’s ρ = −0.76, P = 0.0002) between asynapsis rate and mean expected number of symmetric DSBs (29) based on the chromosome-wide distribution of SPO11 oligos in fertile B6 males (30).

### Asynapsed chromosomes are devoid of active euchromatin

The Cot-1 euchromatin RNA (ecRNA) is localized throughout the active interphase chromosome territory *in cis* and does not diffuse into the rest of the nucleus like other types of RNAs. EcRNA is mainly composed of repeat-rich noncoding RNA in active, open euchromatin (Hall *et al.* 2014). Accordingly, we used Cot-1 DNA as a probe for ecRNA FISH to compare the ecRNA distribution in asynapsed and synapsed chromosomes. Pachytene spermatocytes of PB6F1 hybrids were examined for subnuclear localization of active chromatin and asynapsed chromosomes using confocal fluorescence microscopy after Cot-1 RNA FISH and HORMAD2 immunolabeling. Fluorescence signal quantification revealed that subnuclear regions of asynapsed chromosomes composed of sex chromosomes and/or autosomal univalents were lacking active euchromatin in contrast to other regions of the pachytene nuclei (*Figure supplement 1B - video supplement 1*). We propose that the absence of active euchromatin is a consequence of the meiotic synapsis failure of intersubspecific chromosomes, known as meiotic silencing of unsynapsed chromatin (MSUC (Burgoyne *et al.* 2009)), which can act as an epigenic component contributing to the meiotic phenotypes of sterile hybrids. Our findings are in agreement with the transcriptional under-expression of genes on several autosomes and disruption of meiotic sex chromosome inactivation (MSCI) in leptotene/zygotene spermatocytes from semi-sterile *Mmm* × *Mmd* male hybrids with *Prdm9^PWK/LEWES^* and *Hstx2^PWK^* or *Hstx2^LEWES^* allelic combinations (Larson *et al.* 2016).

### The minimal length of consubspecific sequence necessary to rescue meiotic chromosome synapsis

We have shown previously that meiotic asynapsis affects intersubspecific (PWD/B6) but not consubspecific (PWD/PWD) pairs of homologous chromosomes in sterile male hybrids from crosses of PWD females and B6.PWD-Chr # consomic males (Gregorova *et al.* 2008; Bhattacharyya *et al.* 2013). Here, we searched for the minimum length of the PWD/PWD consubspecific sequence that still could secure synapsis of a chromosome and potentially restore fertility in the hybrids. Instead of substituting the whole PWD chromosome for its B6 homolog, we generated recombinant PWD/B6 and B6/PWD (centromere/telomere) chromosomes. To do that, we crossed the male hybrids between two B6.PWD-Chr# consomic strains and a PWD female to match the intact maternal PWD homologs and estimated the minimum size and location of consubspecific PWD/PWD stretches needed for synapsis rescue as shown in *Figure 2A*. In three such generated ‘two-chromosome crosses’ (hereafter 2-chr cross) we investigated the effect of the PWD/PWD consubspecific intervals on the asynapsis rate in six different chromosomes - two in a given experiment, namely Chr 5 & Chr 12 (*Figure 2B - table supplement 3 and 4*), Chr 7 & Chr 15 (*Figure 2B - table supplement 5 and 6*) and Chr 17 & Chr 18 (*Figure 2B - table supplement 7 and 8*). Altogether over 12,000 pachynemas from 122 chromosomes were examined. All male progeny of the 2-chr crosses were fully sterile, with low testis weight and the absence of sperm in the epididymis. The analysis of data from six recombinant chromosomes revealed the following common features:

**Figure 2.**
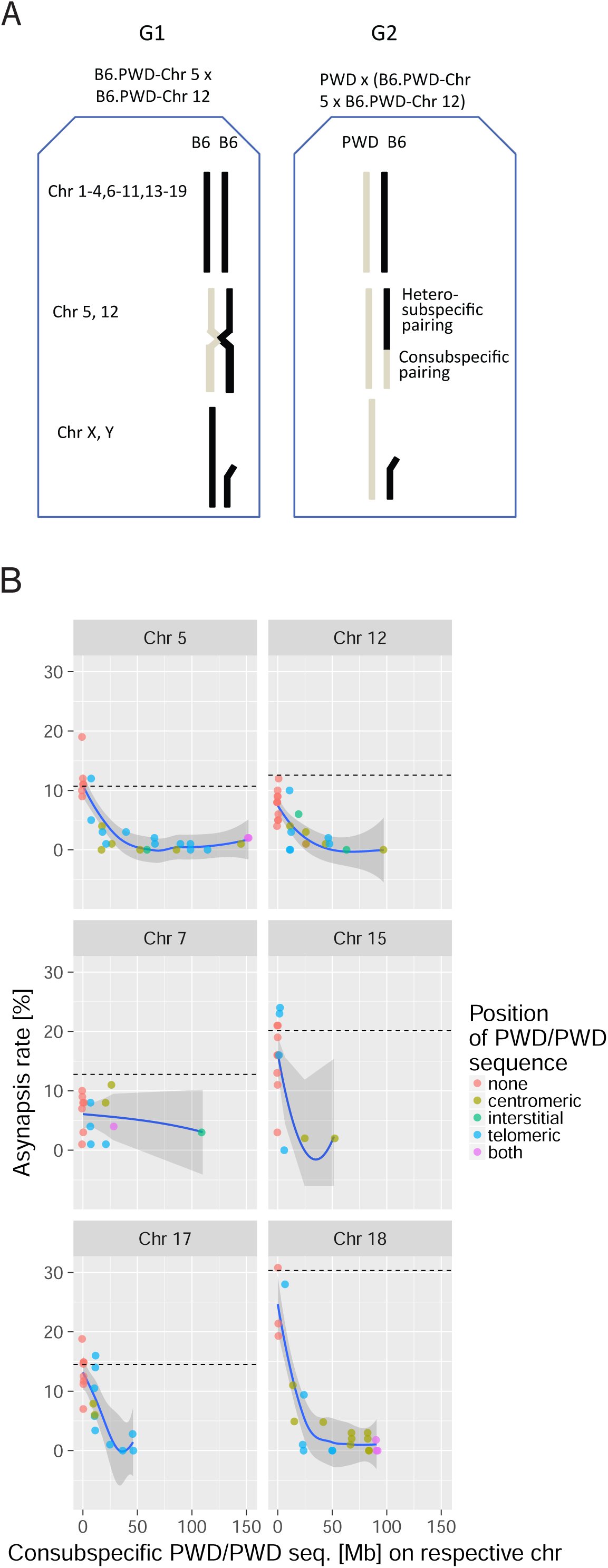
The effect of consubspecific PWD/PWD stretches of genomic sequence on pachytene synapsis, 2-chr cross. (*A*) The F1 hybrid males of two consomic strains (generation 1 - G1, Chr 5 and Chr 12 shown here) were crossed to PWD females to produce generation 2 - G2 sterile F1 hybrids with random recombinant consomic chromosomes 5 and 12. Using whole chromosome probes, the asynapsis rate of the consomic chromosomes was scored by DNA FISH. (*B*) Combination of two chromosomes (5+12, 7+15 and 17+18) were challenged in each experiment. The localization of PWD homozygous sequence with respect to centromere, interstitial part of the chromosome, telomere, or on both ends is distinguished by color (see also Tables Supplement 3 – Supplement 8). The average length between the minimum and maximum of the consubspecific sequence is plotted. The mean asynapsis rate of a given chromosome is regularly higher in PB6F1 hybrids (dashed line) than in 2-chr cross. For explanation see Fig. 4 and the chapter on the transeffect dependent variation in asynapsis rate. Loess curve with 95% CI.

- Introduction by recombination of 27 Mb or more of a consubspecific (PWD/PWD) interval into a pair of intersubspecific (PWD/B6) homologs effectively suppressed the asynapsis rate below the baseline of 5 % in all six studied autosomes (*Figure 2B*). The efficiency of synapsis rescue was gradual with an apparent change point (*Figure 2B*). To describe the pattern in the data a segmented regression model was used (see Materials and Methods). The model based on the data pooled from all 2-chr crosses was selected as the best model with an estimated change point estimate at 27.14 Mb (19.36; 34.91) (95% CI) (see *table supplement 9*). The slope of the decrease of asynapsis in the region of consubspecific intervals shorter than 27.14 Mb Was different for the respective chromosomes, reflecting the asynapsis rate in respective chromosomes in F1 cross (P < 3 χ 10^−11^ F-test).

- In spite of the known role of subtelomeric (bouquet) association in chromosome pairing (Ishiguro *et al.* 2014; Scherthan *et al.* 2014), the location of the consubspecific sequence at the telomeric end was not essential for synapsis. The PWD/PWD intervals of sufficient size located at the centromeric (proximal), interstitial, or telomeric (distal) positions rescued synapsis as well (*Figure 2B - table supplements 3-8*).

### Reversal of hybrid sterility by targeted suppression of asynapsis of four of the most asynapsis-sensitive autosomes

The above experiments have shown that a randomly located consubspecific PWD/PWD interval with a 27 or more Mb on otherwise intersubspecific PWD/B6 background is sufficient to restore the pachytene synapsis of a given autosomal pair. To check the causal relationship between meiotic chromosome asynapsis and HS, we attempted to reverse HS by reducing the asynapsis in the four most asynapsis-prone chromosomes. Provided that hybrid male sterility is directly dependent on chromosome synapsis we predicted, by multiplying the probabilities of the synapsis of individual chromosomes obtained in F1 hybrids, that complete elimination of asynapsis of four of the shortest autosomes (excluding Chr 17 to avoid *Prdm9^PWD/PWD^* interference) could increase the proportion of primary spermatocytes with the full set of synapsed autosomes up to 26.7% and potentially abolish apoptosis of these cells to yield around 5 million sperm cells in the epididymis of the hybrid males.

To evaluate the above prediction experimentally, random intervals of consomic Chrs 15^PWD^, 16 ^PWD^, 18 ^PWD^, and 19 ^PWD^ were transferred on the genetic background of B6 mice in a three-generation cross as shown in *Figure 3A*. Eleven G3 males selected for maximal extent of PWD sequence on these chromosomes were crossed to PWD females (*Figure 3A -table supplement 10*). The resulting G4 hybrid male progeny (hereafter 4-chr cross) displayed various degrees of PWD homozygosity in the studied consomic autosomes on otherwise intersubspecific PWD/B6 genetic background. As predicted, a significant fraction of hybrid males indeed showed partial rescue of spermatogenesis. While in the PB6F1 cross, 100 % of hybrid males displayed no sperm in the epididymis, in the 4-chr cross, only 51.7 % of 87 G4 males were aspermic, 19.5 % had a 0.01–0.74 × 10^6^ sperm count, and 28.7 % had 1.0–13.7 × 10^6^ sperm cells (*Figure 3B - figure supplement 2, table supplement 11*).

**Figure 3.**
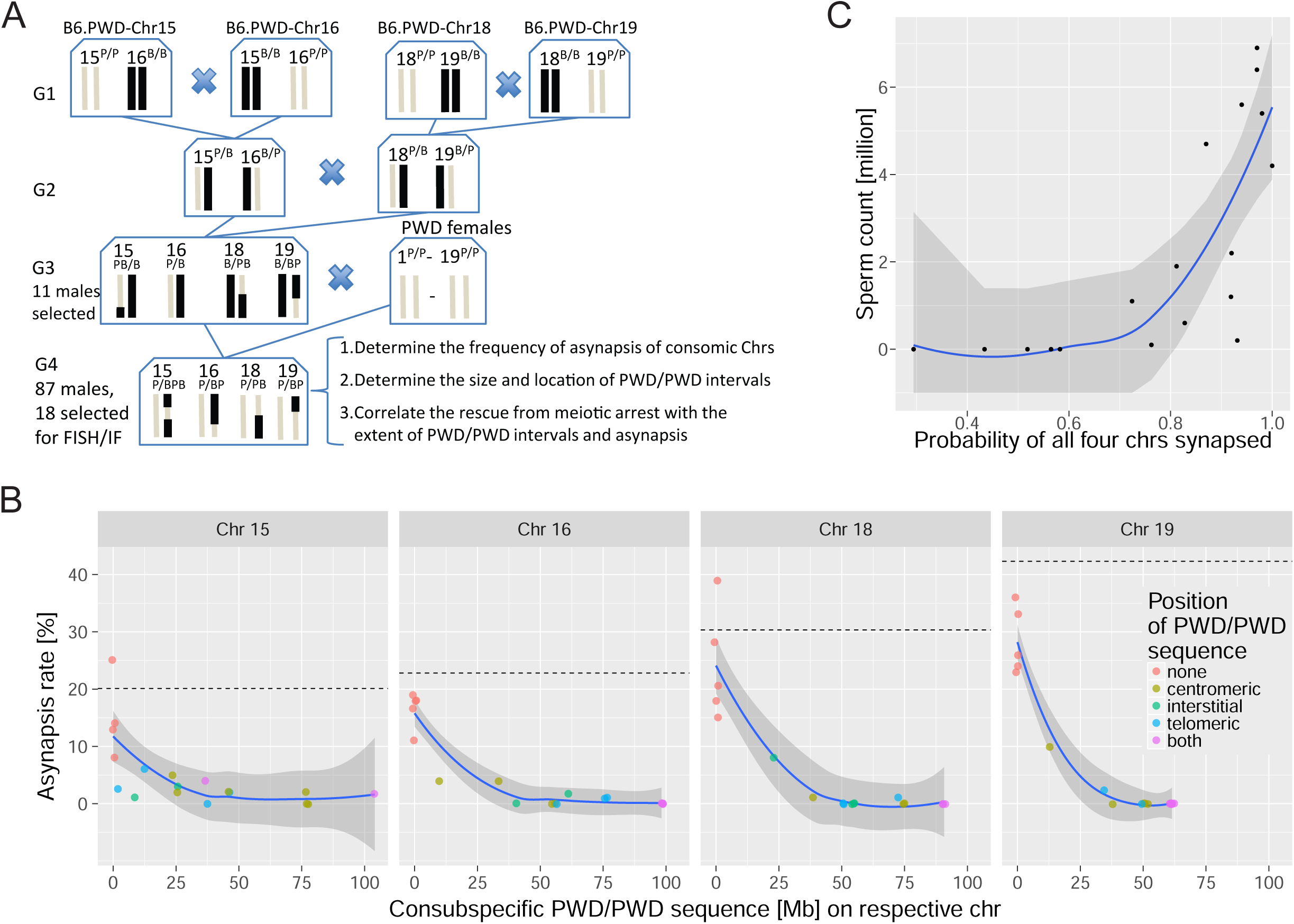
The effect of consubspecific PWD/PWD stretches of genomic sequence on pachytene synapsis and meiotic progression, 4-chr cross. (*A*) Scheme of a four-generation cross resulting in F1 hybrids with four recombinant consomic chromosomes. (*B*) The asynapsis rate related to the size and chromosomal position of the consubspecific PWD/PWD sequence in four consomic chromosomes (15, 16, 18 and 19, see also Table 1). The localization of PWD homozygous sequence with respect to centromere, interstitial part of the chromosome, telomere, or on both ends is distinguished by color (see also Tables 1). (*C*) Number of sperm in epididymis is a function of probability of synapsis of all four consomic chromosomes. The complete meiotic arrest is reversed in males having 70% or higher chance of all four chromosomes synapsed. See also Table Supplement 11. Loess curve with 95% CI.

Next, we asked whether the reversal of meiotic arrest correlates with the recovery of meiotic synapsis of recombined chromosomes and with the size of PWD/PWD consubspecific stretches in the four manipulated chromosomes. Eighteen G4 males were deliberately selected according to their fertility parameters, 13 with HS partial rescue, displaying sperm cells in the epididymis (0,1 × 10^6^−6,9 × 10^6^), and five aspermic controls. The meiotic analysis of over 6500 pachynemas from the genotyped males indeed confirmed the prediction based on the results of 2-chr crosses. The nonrecombinant PWD/PWD consubspecific bivalents were always fully synapsed, while all nonrecombinant PWD/B6 intersubspecific pairs revealed the highest frequencies of asynapsis. All recombinant chromosomes with consubspecific intervals of sufficient length (*Figure 3B -table supplement 12;* see *table supplement 9* for change point estimates) effectively restored synapsis. Moreover, the presence of sperm cells corresponded with the rescue of synapsis of consomic chromosomes. As a rule of thumb, the hybrids had sperm when asynapsis was suppressed in at least three of four segregating chromosomes and when the probability of all four consomic chromosomes being synapsed was > 0.7. Chrs 16, 18, and 19 contributed the strongest effect. (*Figure 3C - table supplement 12*).

### Evidence for a trans-effect on the rate of asynapsis

Provided the probability of failure of the synapsis of each chromosome was completely independent of the rest of the hybrid genome, the asynapsis rate of the nonrecombinant intersubspecific chromosome pairs would be the same in F1 hybrids, 2-chr crosses, and 4-chr cross. Moreover, the frequency of pachynemas with all chromosomes synapsed could be predicted from the observed frequencies of the synapsis of individual chromosomes. Such predicted values would be close to the values directly read from the meiotic spreads and would lie along the diagonal in *Figure 4* As shown below, both types of analysis clearly revealed that the asynapsis rate of a particular chromosome depends on the synapsis status of other chromosomes. In PB6F1 hybrids, the observed 13.1 % (11.4 %, 14.9 %) (95 % CI) of fully synapsed pachynemas was double (P = 0.023, Mann-Whitney test) the rate expected by the multiplication of the observed synapsis rates 6.6 % (5.3 %, 8.1 %) of individual chromosomes (*Figure 4*) indicating a *trans* effect of synapsed autosomes on the probability of the asynapsis of other intersubspecific chromosome pairs. The tendency is more pronounced in data from 2-chr crosses and 4-chr cross experiments. The most straightforward comparison is between the nonrecombinant PWD/B6 intersubspecific chromosomes from F1 hybrids and 2-chr cross or 4-chr crosses, where the asynapsis rate is dramatically reduced to 58-91 % of the respective F1 hybrids’ rates. Moreover, the difference between the observed and expected ratios of asynaptic pachynemas increased with increasing probability of sperm cells in the epididymis (*Figure 4 - table supplement 13*). In 4-chr cross, the *trans* effect was analyzed further by comparing the asynapsis rate of a given nonrecombinant PWD/B6 pair with the other three analyzed chromosomes. *Figure supplement 3* shows a negative correlation (from r= −0.45 for Chr 16 to r = −0.88 for Chr 15, linear regression slope is β_3_chrs_syn_ = −0.19, P = 0.0003) between the asynapsis rate of the nonrecombinant chromosome and the probability of synapsis of the other three studied chromosomes.

**Figure 4.**
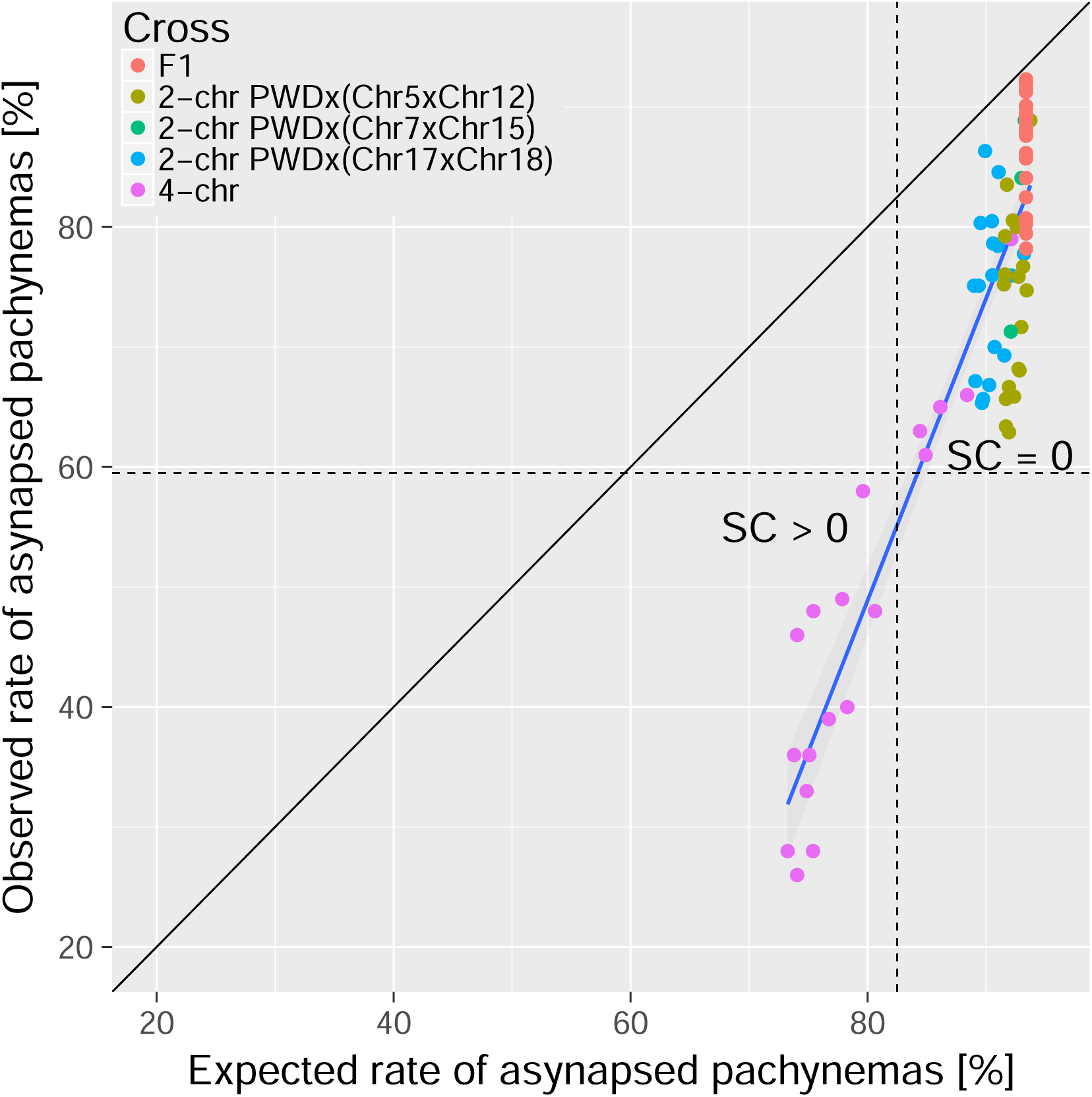
The trans-acting effect of consubspecific PWD/PWD stretches increases the probability of full synapsis of PWD/B6 heterosubspecific homologs in males 2-chr cross and 4-chr cross experiments. The expected rate of synapsed pachynemas was calculated for each mouse in 2-chr cross and 4-chr cross experiments by multiplication of observed synapsis rates (i.e. assuming i ndependence) of FISH analyzed chromosomes (e.g. chrs 15, 16, 18 and 19 in 4-chr cross) and observed PB6F1 synapsis rates of the remaining autosomes. Asynapsis rate was calculated as a complement to synapsis rate. The difference between expected and observed overall asynapsis is most pronounced in 4-chr cross males with the lowest expected overall asynapsis rate. Recovery of spermatogenesis signaled by the presence of sperm in the epididymis occurs when more than 40% of pachynemas are fully synapsed. SC is sperm count.

It has to be emphasized that in the animals with identical *Prdm9* and *Hstx2* and mostly intersubspecific chromosomes investigated in this work, the *cis*-effect of the PWD/PWD length in a given chromosome dominates over the *trans*-effect. Namely, the *trans*-effect can be more clearly observed when conditioned on the *cis*-effect of the PWD/PWD length in a given chromosome (P = 0.002, F-test), while it is harder to detect (P = 0.042, F-test) when the PWD/PWD length in a given chromosome is not taken into account. On the other hand, the asynapsis rate critically depends on the *cis*-effect of the PWD/PWD length in a given chromosome (P < 2×10^−16^, F-test) when the *trans*-effect is not taken into account.

To conclude, the described *trans* effect enhances the probability of the successful pairing of intersubspecific chromosomes in males with identical *Prdm9* and *Hstx2*. The mechanism is unknown, but it is tempting to speculate that a rate-limiting step of an anti-recombination mismatch repair machinery (Spies and Fishel 2015; Chakraborty and Alani 2016) could be involved as elaborated in Discussion.

## Discussion

While the genic control of HS and meiotic synapsis in PBF1 hybrids can be demonstrated by complete restitution of fertility and meiotic pairing in males with *Prdm9^PWD/PWD^* or *Prdm9^PWD/B6Hu^* genotypes and partial recovery in *Prdm9^PWD/C3H^* males (Dzur-Gejdosova *et al.* 2012; Bhattacharyya *et al.* 2013; Davies *et al.* 2016), a chromosome-autonomous nature of asynapsis became apparent in experiments where PB6F1 hybrids carried a single pair of PWD/PWD consubspecific homologs. The males remained sterile, but synapsis of the particular consubspecific pair was completely restored (Bhattacharyya *et al.* 2013). Such regulation of meiotic asynapsis in PBF1 hybrids can be explained by a combined effect of the chromosome-autonomous interaction of homologs operating *in cis* and *Prdm9/Hstx2* incompatibility operating *in trans*. Here we separated the non-genic chromosome autonomous from genic control mechanisms by keeping the sterility-determining allelic combination of the *Prdm9^PWD^*/*Prdm9^B6^* gene and *Hstx2^PWD^* locus constant in all crosses, while successively introgressing stretches of the PWD/PWD consubspecific sequence into eight PWD/B6 intersubspecific autosomal pairs.

### The meiotic asynapsis rate correlates with the presumed paucity of symmetric DSBs in individual chromosomes in sterile hybrids

Davies and coworkers (Davies *et al.* 2016) found that the DNA-binding zinc finger domain of PRDM9 molecule is responsible for sterility in PB6F1 hybrids. In the sterile hybrids, most PRDM9^PWD^-specific hotspots reside on B6 chromosomes and, *vice versa*, most of the PRDM9^B6^ binding sites are activated on PWD chromosomes. This asymmetry could be explained in part by erosion of the PRDM9 binding sites due to preferential transmission to progeny of the mutated hotspots motifs (Boulton *et al.* 1997; Myers *et al.* 2010). In a parallel study, (Smagulova *et al.* 2016) authors identified a novel class of strong hotspots absent in PWD and B6 parents apparently related to asymmetric hotspots described in (Davies *et al.* 2016). Moreover, *Prdm9*-independent ‘default’ hotspots were particularly enriched in Chr X and we noticed that percentage of these ‘default’ hotspots in autosomes correlates with the present data on asynapsis rate in F1 hybrids (Spearman’ s *ρ* = 0.69, P = 0.0012). We assume that these *Prdm9*-independent hotspots could represent the late-forming DSBs on unsynapsed chromosomes and, as such, they may be a consequence rather than the cause of meiotic asynapsis (see (Kauppi *et al.* 2013).

We found that meiotic asynapsis affects each autosomal pair in PB6F1 intersubspecific hybrids at distinctively unequal rates, with shorter chromosomes affected more often than the longer ones. A similar pattern of higher sensitivity of smaller autosomes to the synapsis failure was observed in mice with lowered dosages of SPO11 (Kauppi *et al.* 2013) and consequent twofold DSB reduction. The fact that the asynapsis rate of sterile F1 hybrids correlates better with SPO11-oligo derived DSB density (inferred from B6 mouse strain data (Lange *et al.* 2016) than with the chromosome length brings the first experimental evidence for the idea (Davies *et al.* 2016) linking the asynapsis in sterile PB6F1 hybrids to insufficient number of symmetric DSB hotspots.

### Small stretches of consubspecific sequence restore the synapsis of intersubspecific chromosomes

Provided that a shortage of symmetric DSB hotspots (Davies *et al.* 2016) is the ultimate cause of failure of meiotic synapsis of intersubspecific homologs, then by exchanging the asymmetric hotspots for the symmetric ones the full synapsis could be restored. To experimentally test this prediction, we constructed pairs of PWD/B6 intersubspecific homologs with stretches of PWD/PWD consubspecific intervals, which by definition cannot carry asymmetric hotspots. We found that chromosomes with 27 Mb or longer stretches of consubspecific PWD/PWD sequence always rescued full synapsis in male hybrids between PWD females and consomic B6.PWD-Chr # males. The position of the consubspecific interval along the chromosome was not critical for synapsis rescue, in accordance with the finding that synaptonemal complexes nucleate at multiple recombination sites in each chromosome (Zickler and Kleckner 2015; Finsterbusch *et al.* 2016). We assume that the presence of symmetric DSB hotspots within the PWD/PWD homozygous stretches exceeded the threshold of a minimum number of symmetric DSBs thus rescuing normal meiotic synapsis.

Allowing for the assumptions enumerated in in the first section of Results the number of DSBs necessary for proper synapsis of a given chromosome can be estimated based on the expected distribution of symmetric DSB hotspots on all autosomes and their asynapsis ratios in sterile F1 hybrids (*Table supplement 2*). We aimed to model how the induction and repair of DSBs influence proper meiotic synapsis, and tried to estimate the minimum number of symmetric DSBs per chromosome sufficient for full meiotic synapsis. Our model predicts that in approximately 25 % of cases, a chromosome is asynapsed because there is no symmetric DSB (median of P(asynapsis) / P(0 symmetric DSBs) = 4.5). Assuming a critical threshold of the required DSBs, the remaining 75 % of asynapsis could occur on chromosomes with one symmetric DSB. It is consistent with our data that all asynapsis occurs when there is 0 or 1 symmetric DSBs (median of P(asynapsis) / P(0 or symmetric DSBs) = 1.0) and that two symmetric DSBs per chromosome could be sufficient for full development of the synaptonemal complex, as shown in *Figure 5* for PB6F1 hybrids. The same conclusion also holds true for 2-chr cross and 4-chr cross experiments (*Figure 5 - figure supplement 4 and 5*). The deviations from the diagonal in the *Figure 5* depicting 4-chr cross can be explained by the *trans*-effect described in the Results section.

**Figure 5.**
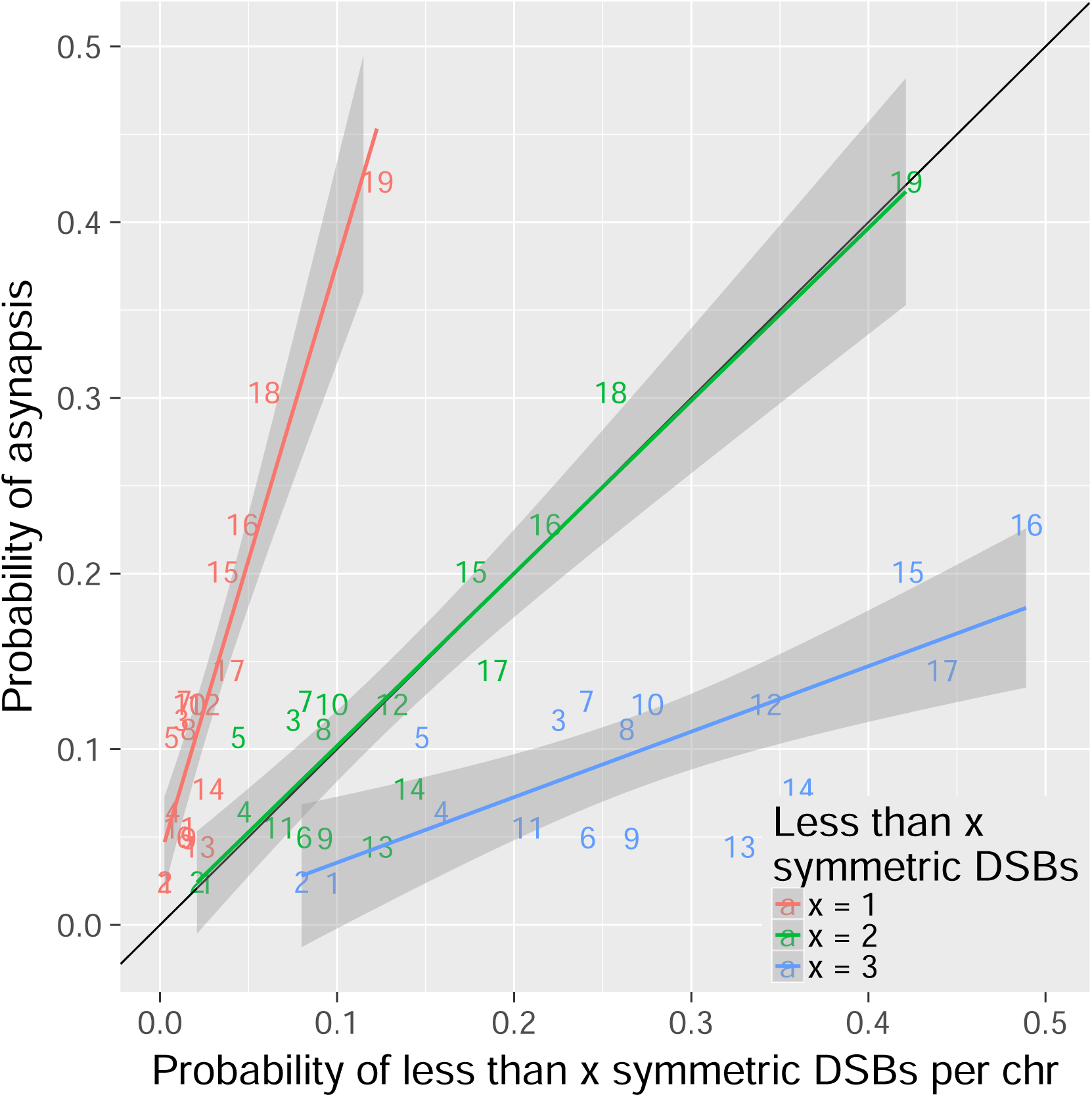
Two or more symmetric DSBs can be sufficient for synapsis. The probability of less than 1 symmetric DSBs per chromosome is ~4 times lower than the asynapsis rate observed in PB6F1 hybrid males, (i.e. estimate of probability of asynapsis) implying ~75% of all asynapsis occurs when there is 1 or more repairable DSBs. The probability of less than 2 symmetric DSBs is a good estimate of the probability of asynapsis, while the probability of less than 3 symmetric DSBs overestimates the probability of asynapsis. This shows that in the simplest explanation two or more symmetric DSBs could be sufficient for synapsis. The probabilistic distribution of symmetric DSBs is calculated based on the model described in the Discussion.

### On the chromosomal nature of hybrid sterility

It has been tacitly assumed, that the early meiotic arrest in sterile PBF1 males is due to the high incidence of asynapsed chromosomes in zygotene/pachytene spermatocytes of PBF1 hybrid males (Bhattacharyya *et al.* 2013). Here we provided the first experimental evidence on linkage between *Prdm9*-controlled asynapsis and meiotic arrest in PBF1 hybrid males by targeted suppression of asynapsis rate in selected autosomes. The suppression of asynapsis in any of three combinations of two PWD/B6 recombinant chromosomes (2-chr cross experiment) did not improve the spermatogenic arrest. However, when hybrid genomes with four short recombinant chromosomes (that are most sensitive to asynapsis) were constructed, almost 50 % of males had sperm in the epididymis and normal testis size. By improving synapsis in three of the four manipulated chromosomes (representing ~11 % of the genome) with randomly located PWD/PWD homozygous intervals, more than 40 % of pachynemas regained full chromosome synapsis and yielded sperm. These results show that the high incidence of primary spermatocytes with asynaptic chromosomes is indeed the factor determining the extent of meiotic arrest in intersubspecific PBF1 hybrids.

The specific feature of PBF1 hybrid sterility consists in the inter-homolog interactions controlled by the *Prdm9* and *Hstx2* genes. The interhomolog incompatibility caused by single nucleotide variation in PRDM9 binding sites, which are situated away from coding genes (Baudat *et al.* 2013) is proposed to make the asymmetric sites difficult to repair (Smagulova *et al.* 2016). We hypothesize that building the meiotic barrier between (sub)species begins with genome-wide accumulation of single nucleotide variants within PRDM9 binding sites and leading to gene conversion-guided meiotic drive (Baker *et al.* 2015). In the subspecific hybrids the asymmetric binding sites can be more difficult to repair using the homolog from the other subspecies as a template because of the strict requirements for sequence identity resulting in the rejection of mismatched heteroduplex intermediates as demonstrated in yeast (Sugawara *et al.* 2004). Originally, such an inter-species barrier was proposed by Radman and colleagues (Rayssiguier *et al.* 1989; Stambuk and Radman 1998) to prevent homeologous recombination between *E. coli* and *Salmonella typhiimurium.* Among eukaryotes, the role of the mismatch repair system in reproductive isolation was reported in *Saccharomyces* species (Hunter *et al.* 1996; Greig *et al.* 2003; Liti *et al.* 2006). An exciting possibility arises that the heteroduplex rejection activity of mismatch repair machinery could operate as a mechanism gradually restricting gene flow between related (sub)species, a *conditio sine qua non* of speciation.

## Materials and Methods

### Mice, Ethics Statement and Genotyping

The mice were maintained at the Institute of Molecular Genetics in Prague and Vestec, Czech Republic. The project was approved by the Animal Care and use Committee of the Institute of Molecular Genetics AS CR, protocol No 141/2012. The principles of laboratory animal care, Czech Act No. 246/1992 Sb., compatible with EU Council Directive 86/609/EEC and Apendix of the Council of Europe Convention ETS, were observed. SSLP markers used for genotyping consomic chromosomes in 2-chr crosses and 4-chr cross are listed in *table supplement 14*. Detailed descriptions of inbred mouse strains, fertility testing and genotyping are given in the *Supplementary text*.

### Antibodies for the immunostaining of spread spermatocytes

Detailed descriptions of the preparation of meiotic spreads and the visualization of SYCP3, _γ_H2AFX, and HORMAD2 proteins on spreads of meiotic cells are given in the *Supplementary Materials and Methods*.

### DNA and RNA FISH

To identify individual chromosomes, XMP Xcyting Mouse Chromosome N Whole Painting Probes (Metasystem) were used for DNA FISH. Identification of nascent RNA in pachytene spermatocytes was performed as previously described (Mahadevaiah *et al.* 2009), with modifications. A detailed description is given in the *Supplementary Materials and Methods*.

### Microscopy and image capture

Images were observed using a Nikon Eclipse 400 epifluorescence microscope and captured with a DS-QiMc mono-chrome camera (Nikon). Images were analyzed using the NIS elements program. A detailed description is given in the *Supplementary Materials and Methods*.

### Statistics

All calculations were performed in the statistical environment R 3.2.2. For modeling the dependence between the asynapsis rate and the number of symmetric DSBs, we determined the probabilistic distribution of the number of symmetric DSBs. The distribution was determined by simulation and with parameters based on previous studies: (i) The number of DSBs per cell (Bhattacharyya *et al.* 2013) was modeled as an observation from the normal distribution N(250, sd=20). (ii) We assumed a number of DSBs proportional to Spo11 oligos (Lange *et al.* 2016) in each autosome. (iii) The positions of DSBs in the particular autosome were simulated from the uniform distribution, U(0, Autosome length). (iv) For the intersubspecific part of the autosome, the number of symmetric DSBs was simulated from the binomial distribution Bi(n=N_DSBs_in_het_part, p=0.28 (Davies *et al.* 2016; Smagulova *et al.* 2016) For the consubspecific part of the autosome, all DSBs were taken as symmetric. The total number of symmetric DSBs in the autosome was taken as the sum of symmetric DSBs in the respective parts.

Steps (i)-(iv) were performed in N=100000 simulations to obtain a probabilistic distribution. Further details on statistical evaluation are given in the *Supplementary Materials and Methods*.

## Acknowledgments

We thank Simon Myers for sharing his unpublished data and critical comments, Attila Tóth for providing the HORMAD2 antibody, Mary Ann Handel and Linda Odenthal-Hesse for critical reading of this manuscript, David Green, Sarka Takacova, and members of the Forejt lab for their helpful comments, M. Capek for help with analysis of confocal microscopy data, and J. Perlova for her assistance with genotyping. This work was supported by Czech Science Foundation grant 13-08078S and by the LQ1604 project of the NSPII from the Ministry of Education, Youth and Sports of the Czech Republic. Barbora Valiskova was partly supported by project GA UK No.17115 from Charles University, Czech Republic. We also acknowledge the Light Microscopy Core Facility, IMG ASCR, Prague, supported by MEYS (LM2015062), OPPK (CZ.2.16/3.1.00/21547) and (LO1419).

## Supplementary Materials and Methods

### Mice, ethics statement

The project was approved by the Animal Care and use Committee of the Institute of Molecular Genetics AS CR, protocol No 141/2012. The principles of laboratory animal care, Czech Act No. 246/1992 Sb., compatible with EU Council Directive 86/609/EEC and Apendix of the Council of Europe Convention ETS, were observed. SSLP markers used for genotyping consomic chromosomes in 2-chr crosses and 4-chr cross are listed in Table supplement 14.

The PWD/Ph inbred strain originated from a single pair of wild mice of the ***Mus musculus musculus*** subspecies trapped in 1972 in Central Bohemia, Czech Republic (Gregorova and Forejt, 2000). The C57BL/6J (B6) inbred strain was purchased from The Jackson Laboratory. The panel of 27 chromosome substitution strains C57BL/6J-Chr #PWD (abbreviated B6.PWD-Chr#) was prepared in our laboratory and is maintained by the Institute of Molecular Genetics AS CR, Division BIOCEV, Vestec, Czech Republic, and by The Jackson laboratory, Bar Harbor, Maine, U.S.A. All mice were maintained in a 12-h light/12-h dark cycle in a specific pathogen-free barrier facility. The mice had ad libitum access to a standard rodent diet (ST-1, 3.4% fat; VELAZ) and water. All males were killed at age 60-70 d.

### Immunostaining and image capture

For immunocytochemistry, the spread nuclei were prepared as described (Anderson et al. 1999) with modifications. Briefly, a single-cell suspension of spermatogenic cells in 0.1M sucrose with Protease inhibitors (Roche) was dropped on 1% paraformaldehyde-treated slides and allowed to settle for 3 hours in a humidified box at 4 °C. After brief H_2_O and PBS washing and blocking with 5% goat sera in PBS (vol/vol), the cells were immunolabeled using a standard protocol with the following antibodies: anti-HORMAD2 (1:700, rabbit polyclonal antibody, gift from Attila Toth) and SYCP3 (1:50, mouse monoclonal antibody, Santa Cruz, #74569). Secondary antibodies were used at 1:500 dilutions and incubated at room temperature for 60 min; goat anti-Rabbit IgG-AlexaFluor568 (MolecularProbes, A-11036) and goat anti-Mouse IgG-AlexaFluor647 (MolecularProbes, A-21235). The images were acquired and examined in a Nikon Eclipse 400 microscope with a motorized stage control using a Plan Fluor objective, 60x (MRH00601; Nikon) and captured using a DS-QiMc monochrome CCD camera (Nikon) and the NIS-Elements program (Nikon). The images were processed using the Adobe Photoshop CS software (Adobe Systems).

### Combined immunofluorescence staining with DNA FISH or RNA FISH

XMP XCyting Mouse Chromosome N Whole Painting Probes (Metasystems) were used for DNA FISH analysis of asynapsis of all autosomes, one at a time, as described (Turner et al. 2005), with slight modifications. Testes from 8-week-old mice were dissected and spread meiocyte nuclei were prepared as described previously (Mahadevaiah et al. 2009) with a modification, which relies on reversed sequence of RNA FISH and immunofluorescence staining (Turner et al. 2005). Briefly, after cell fixation and permeabilization, the immunofluorescent labeling was performed for 90 min at 20°C with primary anti HORMAD2 and anti SYCP3 antibodies. Secondary antibodies were selected as above and incubated at room temperature for 60 min. After washing and postfixation steps, the immunostained nuclei were processed with RNA fluorescence in situ hybridization. The Cot-1 DNA biotin-labeled probe was incubated overnight at 37°C, and then the hybridized biotinylated Cot-1 probe was labelled with FITC-avidin conjugate and the fluorescent signal was amplified as described previously (Chaumeil et al, 2008). The images of the immunofluorescence stained and cot-1 RNA FISH-labeled spread spermatocytes were examined and photographed using confocal microscope DMI6000CEL - xsLeica TCS SP8.

### Statistics

The effects of the number of Spo11 oligos and the chromosomal length on the asynapsis rate were investigated using a GLMM model. In all the used GLMM models in this work, the asynapsis was modeled as a binary response to fixed effects under investigation and a random intercept for each animal. In F1 hybrids 95 % confidence intervals of observed rate of asynapsed pachynemas and expected rated of asynapsed pachynemas were calculated by bootstrap. The estimates of mean asynapsis rate in respective chromosomes, their standard errors and 95 % confidence intervals were based on GLMM model. In 2-chr crosses and 4-chr cross 95 % confidence intervals of asynapsis rate were constructed based on the likelihood ratio to capture also the uncertainty in the cases when zero asynapsis rate per mouse and chromosome was observed.

Based on the nature of the dependence between the asynapsis rate and the length of consubspecific PWD/PWD region on chromosomes of 2-chr crosses and 4-chr cross, we fitted the data by segmented two-part continuous regression models (Vito et al. 2003)

We fitted the models for all the chromosomes separately (Table supplement 9) being aware of the limitations caused by the lack of animals having specific lengths of consubspecific PWD/PWD region in the respective chromosomes. As the best model describing the dependence of asynapsis rate on the lenghts of PWD/PWD intervals we selected piecewise linear models fitting 1) pooled data from 2-chr crosses and 2) pooled data from both 2-chr crosses and 4-chr cross. Those models are not severely affected by the lack of observations with specific lengths of PWD/PWD segment nor by outliers.

All calculations were performed in R 3.2.2, the change point models and GLMM models were fitted using the packages segmented and lme4, respectively (Vito et al. 2008, Douglas et al. 2015).

---

**Figure supplement 1.**
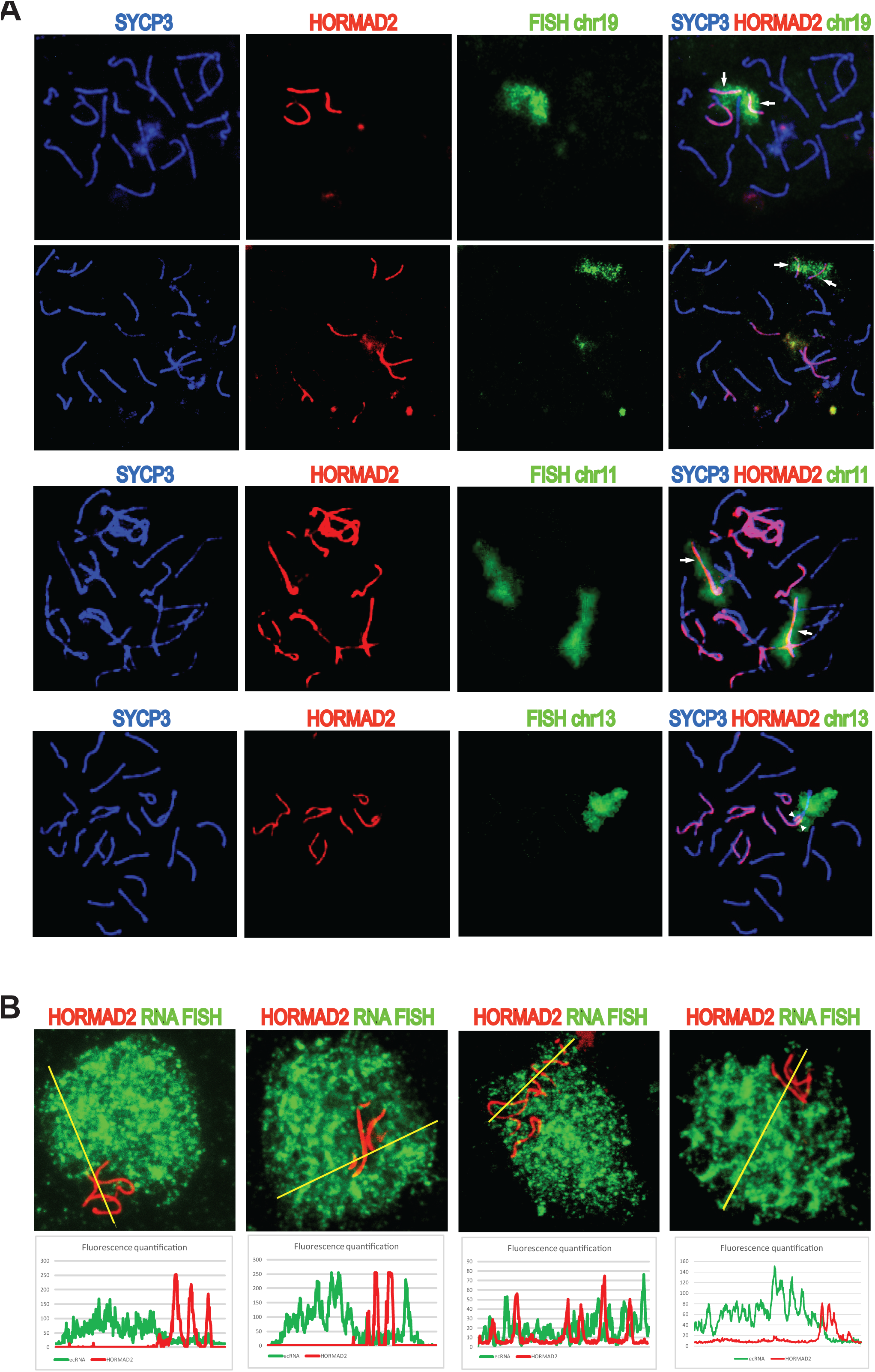
Asynapsis of heterosubspecific homologs in PB6F1 pachynemas. (A) Partial (arrowheads) and complete (arrows) asynapsis of Chr 19, 11 and 13. HORMAD2-labeled chromosomes with synapsis defects often form tangles via nonhomologous pairing. (B) Asynapsed chromosomes are embedded in transcriptionally silenced chromatin visualized by the lack of ecRNA detected by Cot1 RNA FISH See also video 1.

**Figure supplement 2.**
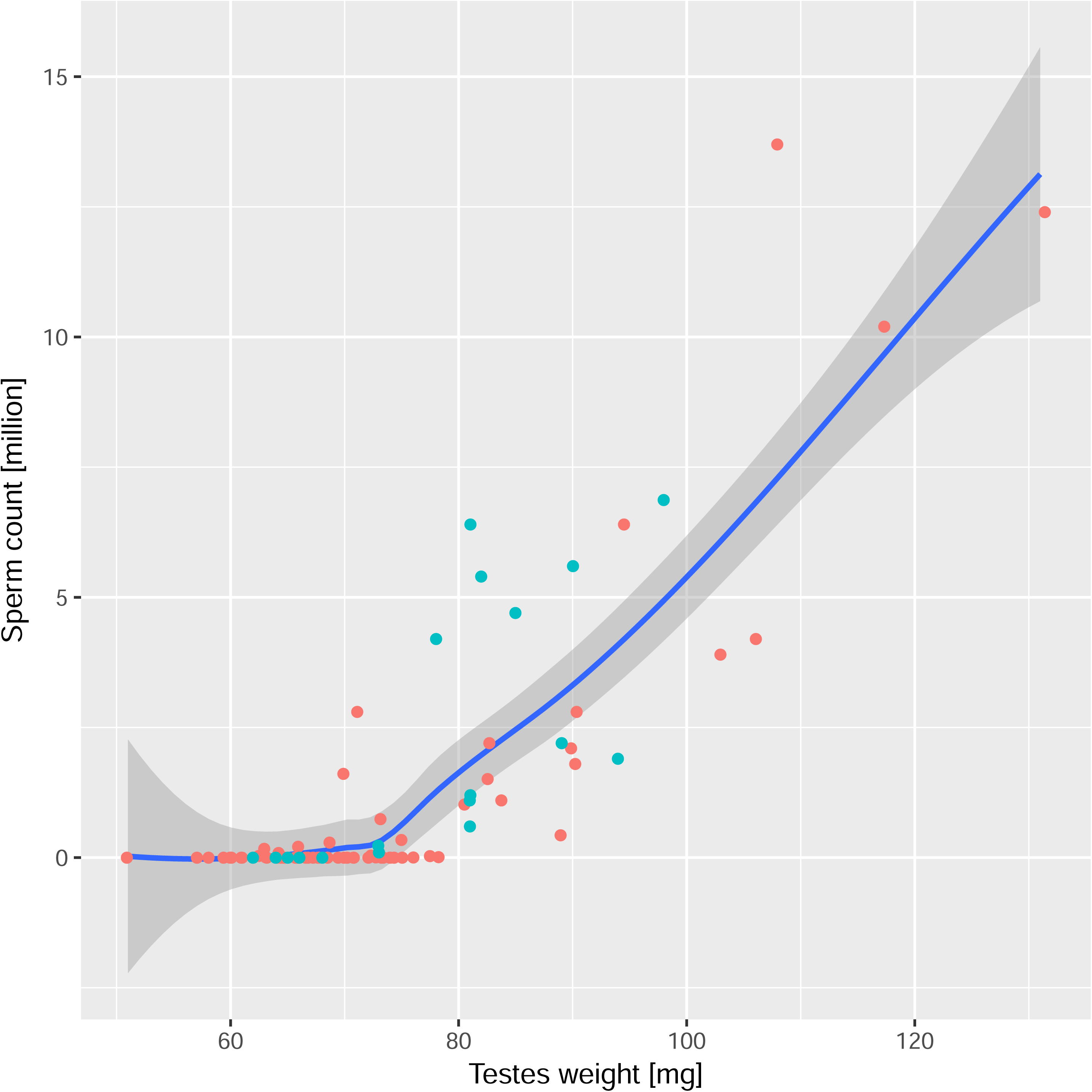
Fertility parameters of G4 males from the 4-chr cross. Rescue of HS meiotic arrest is detectable in males with the weight of paired testes >70 mg and >0.1 × 10^6^ of sperm in the epididymis. The males selected for DNA FISH/HORMAD2 analysis of asynapsis are highlighted in turquoise. All males share the sterility-determining allelic combination of *Prdm9^PWD/B6^* and *Hstx2^PWD^*. Loess curve with 95% CI.

**Figure supplement 3.**
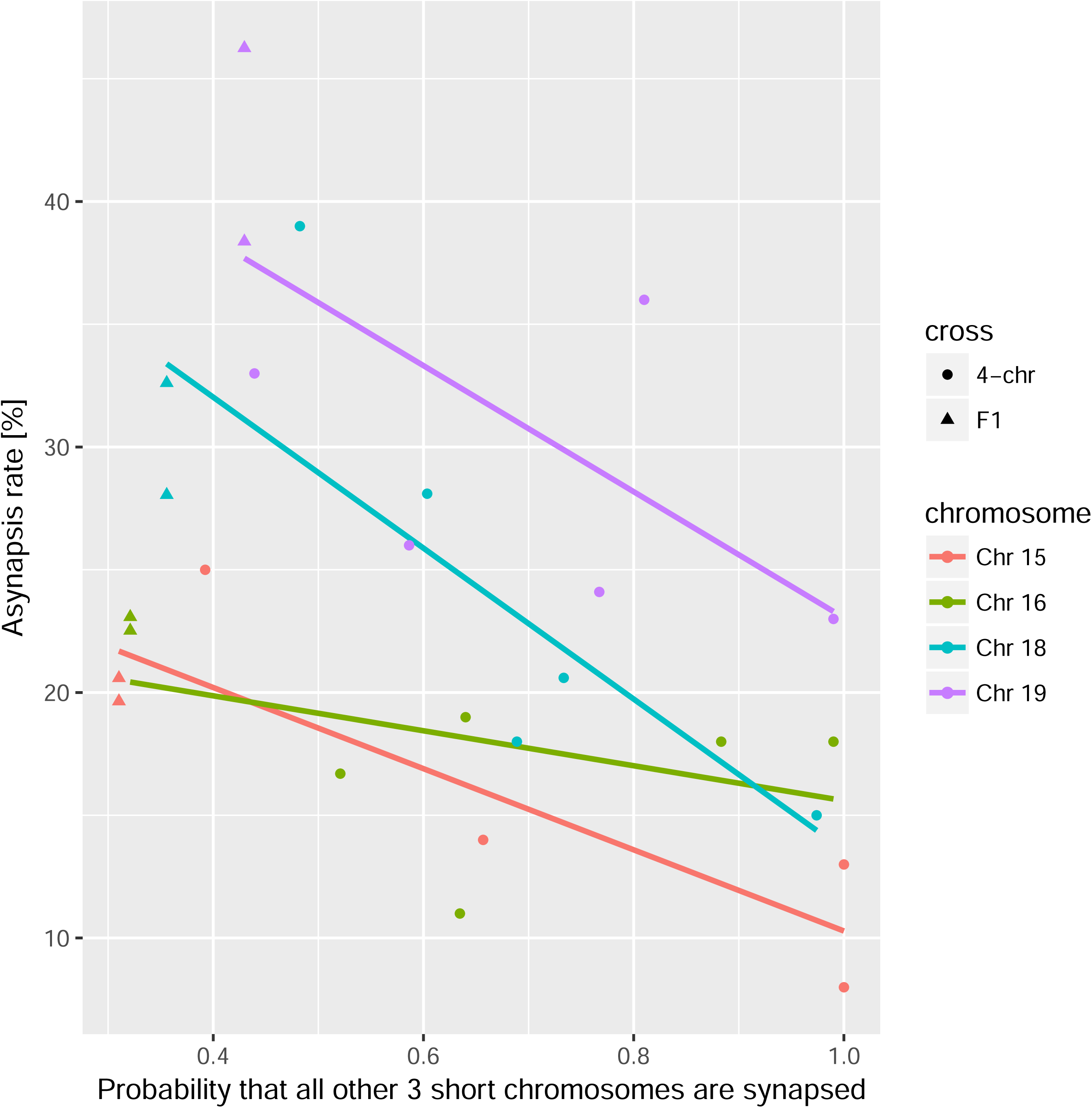
Asynapsis rate of individual nonrecombinant consomic PWD/B6 chromosomes modified in trans by the probability of synapsis of the remaining 3 consomic chromosomes in 4-chr cross in individual males compared to PB6F1 hybrids. Asynapsis rate of a given chromosome is in negative correlation (r = −0.88, −0.45, −0.80, −0.67 for Chrs 15, 16, 18, 19 respectively) with the probability that all other 3 analyzed chromosomes are synapsed. The slope of linear regression is β_3_chrs_syn_ = −0.19, overall P = 0.0003.

**Figure supplement 4.**
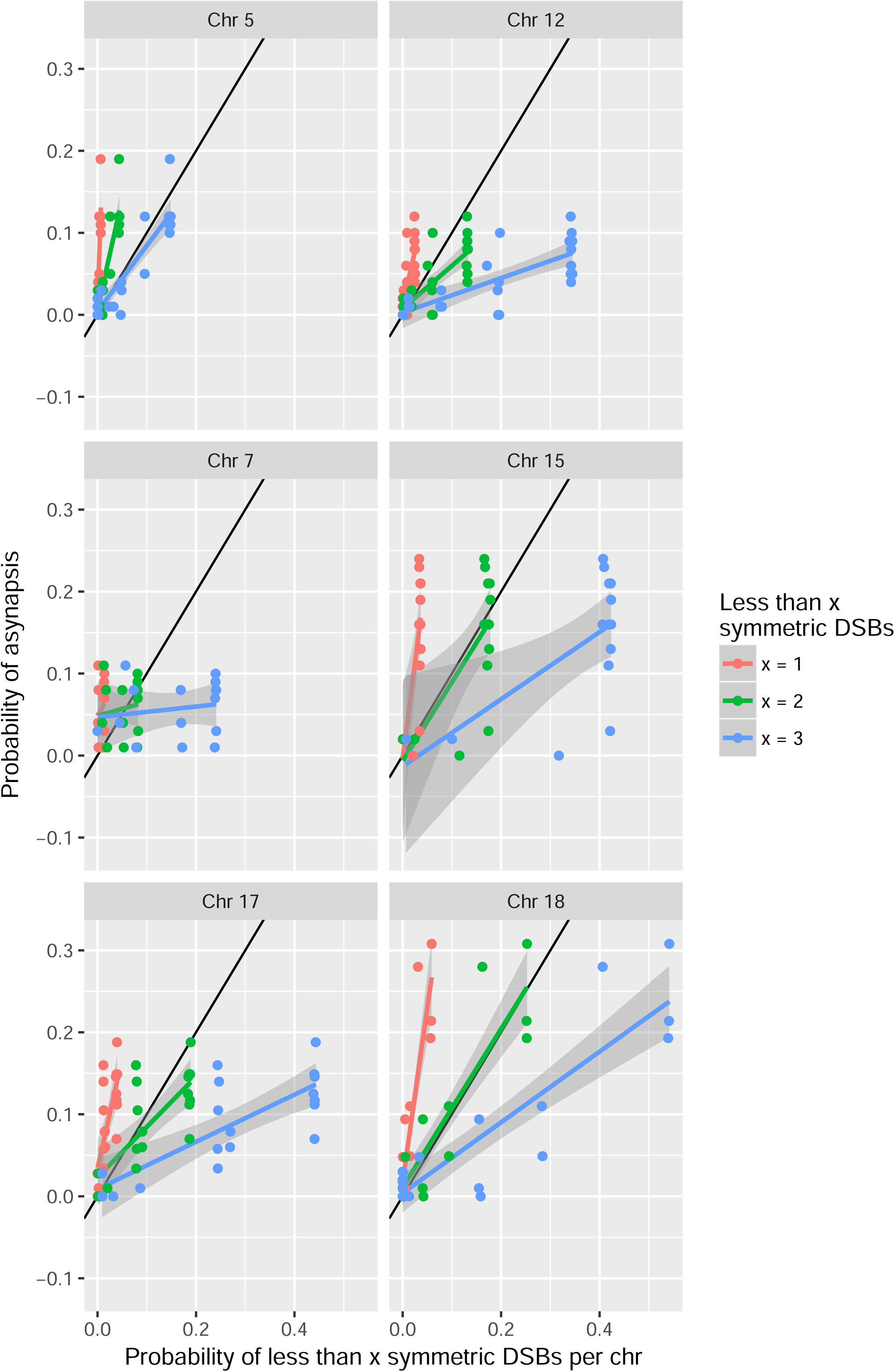
Two and more symmetric DSBs can be sufficient for synapsis in 2-chr cross males. See Fig. 5 for details.

**Figure supplement 5.**
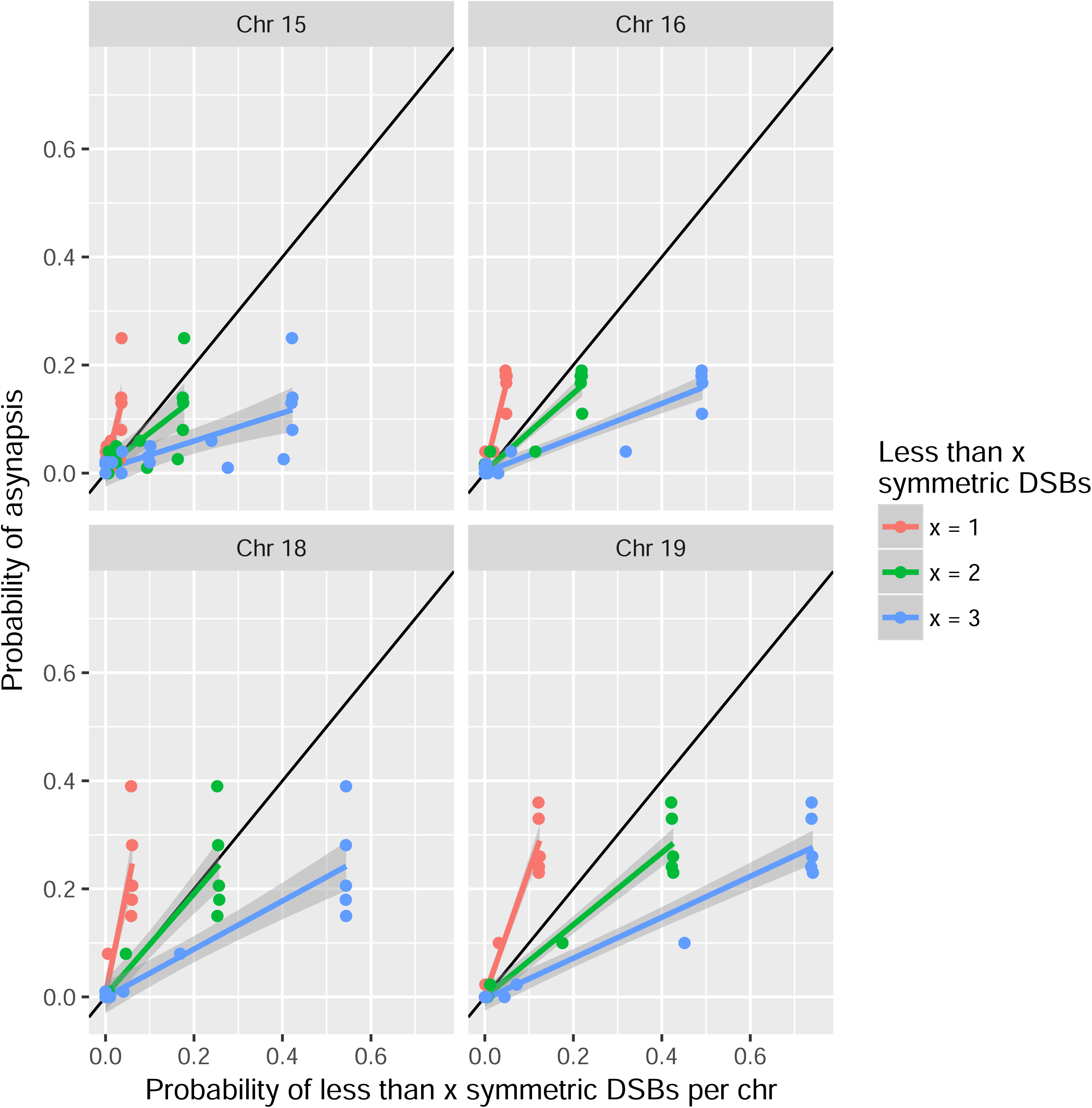
Two or more symmetric DSBs can be sufficient for synapsis in 4-chr cross hybrid males. See Fig. 5 for details.

**Table supplement 1.** Asynapsis rate of individual chromosomes of (PWD x B6)F1 malesand on fertility parameters

| Chromosome | Asynapsis | Asynapsis | Synapsis | Complete Asynapsis | Partial Asynapsis | Intermingled Asynapsis | Number of Males per Chr | Analyzed cells | Complete Asynapsis | Partial Asynapsis | Intermingled Asynapsis |
| --- | --- | --- | --- | --- | --- | --- | --- | --- | --- | --- | --- |
| Name | Mean [%] | 95% CI | Mean [%] | Mean [%] | Mean [%] | Mean [%] | N | n, per male | n, per male | n, per male | n, per male |
| chr1 | 2,6 | 1.2; 5.8 | 97,4 | 0,4 | 2,2 | 0,0 | 2 | 125; 102 | 1; 0 | 5; 0 | 0; 0 |
| chr2 | 2,7 | 1.3; 5.6 | 97,3 | 1,6 | 1,2 | 0,0 | 2 | 103; 152 | 1; 3 | 0; 3 | 0; 0 |
| chr3 | 11,6 | 7.8; 16.8 | 88,4 | 6,5 | 2,5 | 2,5 | 2 | 81; 118 | 4; 9 | 2; 3 | 4; 1 |
| chr4 | 6,5 | 3.9; 10.7 | 93,5 | 0,5 | 4,6 | 1,4 | 2 | 110; 105 | 1; 0 | 7; 3 | 2; 1 |
| chr5 | 10,6 | 7.6; 14.6 | 89,4 | 8,3 | 1,3 | 1,0 | 3 | 85; 89; 129 | 6; 9; 10 | 0; 3; 1 | 0; 2; 1 |
| chr6 | 5,1 | 2.7; 9.6 | 94,9 | 1,1 | 4,0 | 0,0 | 2 | 70; 105 | 1; 1 | 2; 5 | 0; 0 |
| chr7 | 12,8 | 8.8; 18.2 | 87,2 | 8,7 | 4,1 | 0,0 | 2 | 94; 102 | 10; 7 | 2; 6 | 0; 0 |
| chr8 | 10,7 | 7.3; 15.4 | 89,3 | 3,6 | 7,1 | 0,0 | 2 | 93; 132 | 3; 5 | 10; 6 | 0; 0 |
| chr9 | 5,0 | 2.8; 8.8 | 95,0 | 0,5 | 4,5 | 0,0 | 2 | 107; 113 | 0; 1 | 3; 7 | 0; 0 |
| chr10 | 12,6 | 8.8; 17.6 | 87,4 | 4,9 | 5,8 | 1,8 | 2 | 114; 109 | 8; 3 | 3; 10 | 2; 2 |
| chr11 | 5,9 | 3.4; 10.2 | 94,1 | 1,0 | 1,5 | 3,5 | 2 | 126; 76 | 2; 0 | 2; 1 | 5; 2 |
| chr12 | 12,6 | 8.5; 18.2 | 87,4 | 10,4 | 2,2 | 0,0 | 2 | 92; 91 | 10; 9 | 2; 2 | 0; 0 |
| chr13 | 4,4 | 2.6; 7.4 | 95,6 | 3,5 | 1,0 | 0,0 | 3 | 112; 106; 97 | 3; 4; 4 | 2; 0; 1 | 0; 0; 0 |
| chr14 | 7,8 | 5.2; 11.7 | 92,2 | 1,9 | 4,8 | 1,1 | 2 | 129; 139 | 4; 1 | 3; 10 | 0; 3 |
| chr15 | 20,1 | 15.3; 26.0 | 79,9 | 15,0 | 4,7 | 0,5 | 2 | 102; 112 | 18; 14 | 3; 7 | 0; 1 |
| chr16 | 22,8 | 17.5; 29.1 | 77,2 | 15,3 | 6,4 | 1,0 | 2 | 111; 91 | 16; 15 | 7; 6 | 2; 0 |
| chr17 | 14,6 | 10.4; 20.1 | 85,4 | 12,1 | 2,4 | 0,0 | 2 | 111; 95 | 12; 13 | 5; 0 | 0; 0 |
| chr18 | 30,5 | 24.1; 37.7 | 69,5 | 29,9 | 0,6 | 0,0 | 2 | 92; 82 | 29; 23 | 1; 0 | 0; 0 |
| chr19 | 42,2 | 34.9; 49.8 | 57,8 | 39,8 | 2,4 | 0,0 | 2 | 80; 86 | 36; 30 | 1; 3 | 0; 0 |

Synapsis and classes of asynapsis are based on the GLMM model to take into account the unequal numbers of cells analyzed per males.

**Table supplement 3.**
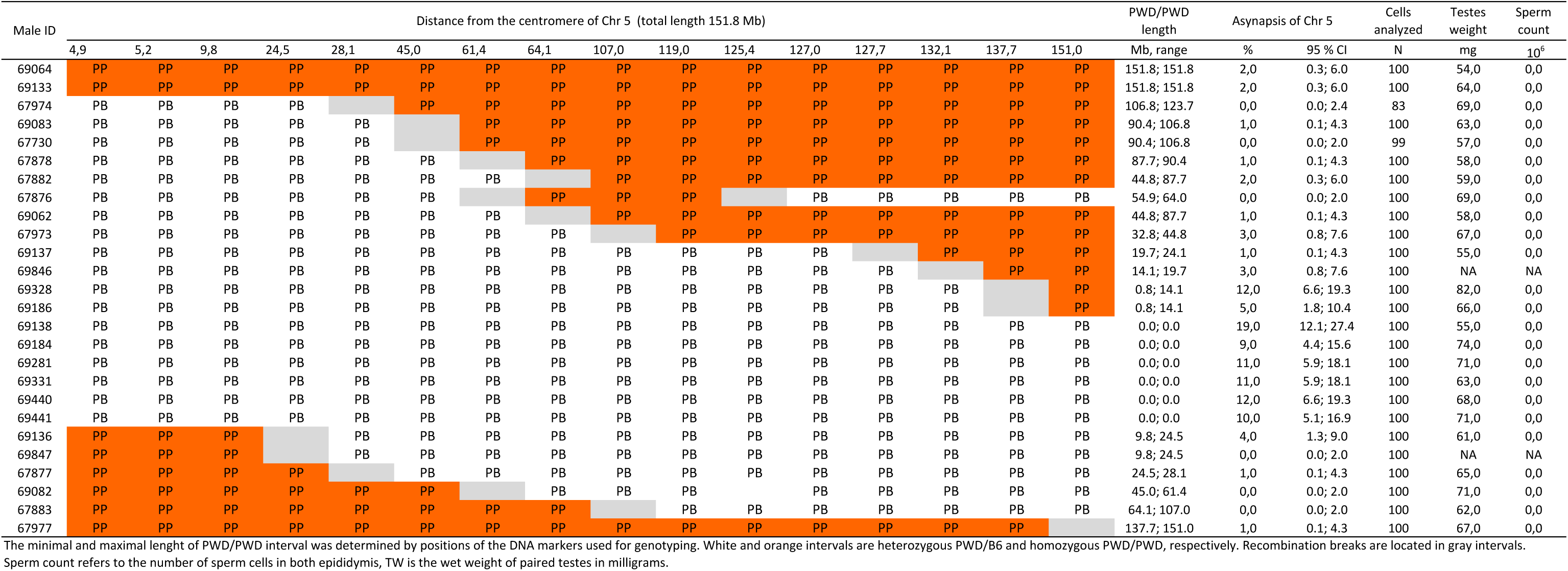
The effect of the size and location of PWD/PWD consubspecific intervals on asynapsis of Chr 5 and on fertility parameters

**Table supplement 4.** The effect of the size and location of PWD/PWD consubspecific intervals on asynapsis of Chr 12 and on fertility parameters

| Male ID | Distance from centromere of Chr 12 (total length 120.1 Mb) |  |  |  |  |  |  |  |  |  | PWD/PWD length | Asynapsis of Chr 12 |  | Cells analyzed | Testes weight | Sperm count |
| --- | --- | --- | --- | --- | --- | --- | --- | --- | --- | --- | --- | --- | --- | --- | --- | --- |
|  | 5,4 | 7,6 | 15,7 | 19,8 | 30,8 | 35,0 | 54,6 | 91,0 | 103,4 | 113,8 | Mb, range | % | 95% CI | N | mg | 10<sup>6</sup> |
| 69064 | PP | PP | PP | PP | PP | PP | PP | PP |  | PB | 91.0; 103.4 | 0,0 | 0.0; 2.0 | 100 | 54,0 | 0,0 |
| 67977 | PB | PB | PB | PB | PB |  | PP | PP | PP |  | 48.8; 78.8 | 0,0 | 0.0; 2.0 | 100 | 67,0 | 0,0 |
| 67876 | PP | PP | PP | PP | PP | PP |  | PB | PB | PB | 35.0; 54.6 | 1,0 | 0.1; 4.3 | 100 | 69,0 | 0,0 |
| 69144 | PP | PP | PP | PP |  | PB | PB |  | PP | PP | 36.5; 59.9 | 1,0 | 0.1; 4.3 | 100 | 66,0 | 0,0 |
| 69760 | PP | PP | PP | PP |  | PB | PB | PB | PB | PB | 19.8; 30.8 | 1,0 | 0.1; 4.3 | 100 | 65,0 | 0,0 |
| 69133 | PP | PP | PP | PP |  | PB | PB | PB | PB | PB | 19.8; 30.8 | 3,0 | 0.8; 7.6 | 100 | 64,0 | 0,0 |
| 67973 |  | PP | PP | PP |  | PB | PB | PB | PB | PB | 12.2; 25.4 | 6,0 | 2.4; 11.8 | 100 | 67,0 | 0,0 |
| 69281 | PP | PP |  | PB | PB | PB | PB | PB | PB | PB | 7.6; 15.7 | 4,0 | 1.3; 9.0 | 100 | 71,0 | 0,0 |
| 67974 | PB | PB | PB | PB | PB | PB | PB | PB | PB | PB | 0.0; 0.0 | 10,0 | 5.1; 16.9 | 100 | 69,0 | 0,0 |
| 69136 | PB | PB | PB | PB | PB | PB | PB | PB | PB | PB | 0.0; 0.0 | 9,0 | 4.4; 15.6 | 100 | 61,0 | 0,0 |
| 69138 | PB | PB | PB | PB | PB | PB | PB | PB | PB | PB | 0.0; 0.0 | 9,0 | 4.4; 15.6 | 100 | 55,0 | 0,0 |
| 69184 | PB | PB | PB | PB | PB | PB | PB | PB | PB | PB | 0.0; 0.0 | 8,0 | 3.7; 14.4 | 100 | 71,0 | 0,0 |
| 69282 | PB | PB | PB | PB | PB | PB | PB | PB | PB | PB | 0.0; 0.0 | 6,0 | 2.4; 11.8 | 100 | 67,0 | 0,0 |
| 69328 | PB | PB | PB | PB | PB | PB | PB | PB | PB | PB | 0.0; 0.0 | 12,0 | 6.6; 19.3 | 100 | 66,0 | 0,0 |
| 69331 | PB | PB | PB | PB | PB | PB | PB | PB | PB | PB | 0.0; 0.0 | 5,0 | 1.8; 10.4 | 100 | 63,0 | 0,0 |
| 69441 | PB | PB | PB | PB | PB | PB | PB | PB | PB | PB | 0.0; 0.0 | 8,0 | 3.7; 14.4 | 100 | 68,0 | 0,0 |
| 69663 | PB | PB | PB | PB | PB | PB | PB | PB | PB | PB | 0.0; 0.0 | 4,0 | 1.3; 9.0 | 100 | 70,0 | 0,0 |
| 69664 | PB | PB | PB | PB | PB | PB | PB | PB | PB | PB | 0.0; 0.0 | 8,0 | 3.7; 14.4 | 100 | 71,0 | 0,0 |
| 69847 | PB | PB | PB | PB | PB | PB | PB | PB | PB | PB | 0.0; 0.0 | 5,0 | 1.8; 10.4 | 100 | 63,0 | 0,0 |
| 69186 | PB | PB | PB | PB | PB | PB | PB | PB |  | PP | 6.3; 16.7 | 10,0 | 5.1; 16.9 | 100 | 82,0 | 0,0 |
| 69763 | PB | PB | PB | PB | PB | PB | PB | PB |  | PP | 6.3; 16.7 | 3,0 | 0.8; 7.6 | 100 | 68,0 | 0,0 |
| 69427 | PB | PB | PB | PB | PB | PB | PB | PB |  | PP | 6.3; 16.7 | 0,0 | 0.0; 2.0 | 100 | 68,0 | 0,0 |
| 69082 | PB | PB | PB | PB | PB | PB | PB | PB |  | PP | 6.3; 16.7 | 0,0 | 0.0; 2.0 | 100 | 71,0 | 0,0 |
| 69062 | PB | PB | PB | PB | PB | PB |  | PP | PP | PP | 29.1; 65.5 | 2,0 | 0.3; 6.0 | 100 | 59,0 | 0,0 |
| 69083 | PB | PB | PB | PB | PB | PB |  | PP | PP | PP | 29.1; 65.5 | 1,0 | 0.1; 4.3 | 100 | 57,0 | 0,0 |

The minimal and maximal length of PWD/PWD interval was determined by positions of the DNA markers used for genotyping. White and orange intervals are heterozygous PWD/B6 and homozygous PWD/PWD, respectively. Recombination breaks are located in gray intervals.
Sperm count refers to the number of sperm cells in both epididymis, TW is the wet weight of paired testes in milligrams.

**Table supplement 5.**
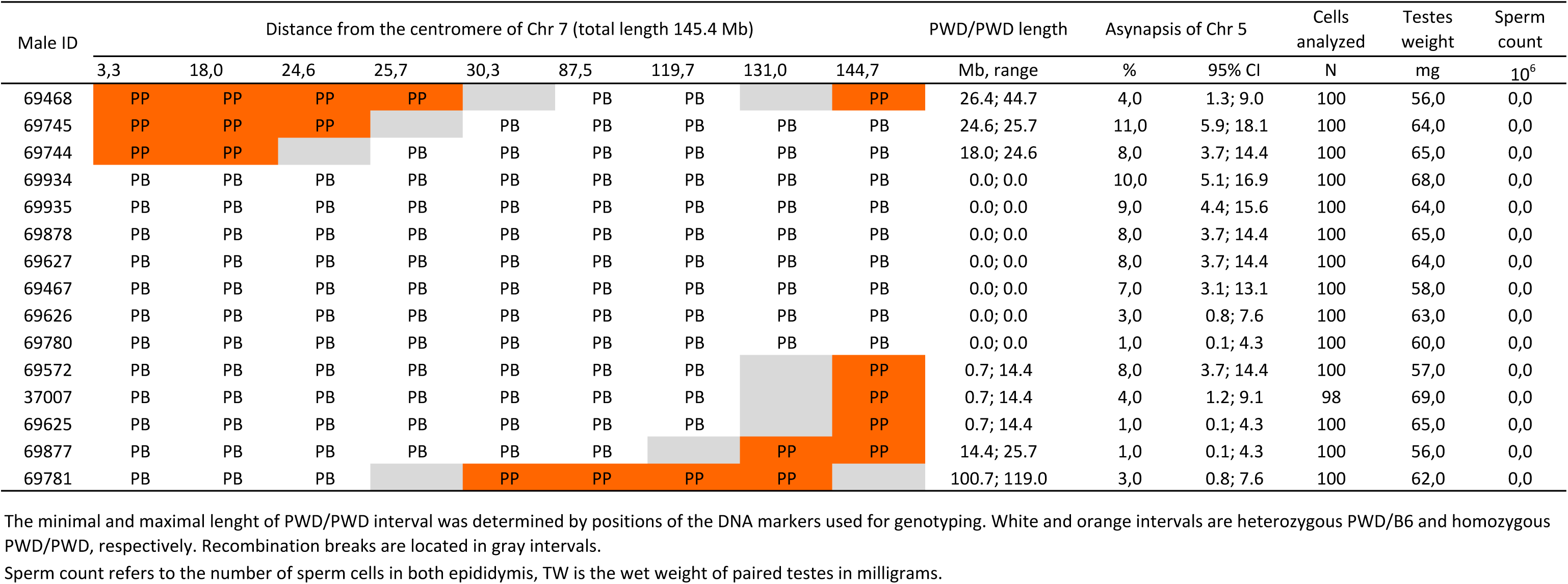
The effect of the size and location of PWD/PWD consubspecific intervals on asynapsis of Chr 7 and on fertility parameters

| Male ID | Distance from the centromere of Chr 7 (total length 145.4 Mb) |  |  |  |  |  |  |  |  | PWD/PWD length | Asynapsis of Chr 5 |  | Cells analyzed | Testes weight | Sperm count |
| --- | --- | --- | --- | --- | --- | --- | --- | --- | --- | --- | --- | --- | --- | --- | --- |
|  | 3,3 | 18,0 | 24,6 | 25,7 | 30,3 | 87,5 | 119,7 | 131,0 | 144,7 | Mb, range | % | 95% CI | N | mg | 10<sup>6</sup> |
| 69468 | PP | PP | PP | PP |  | PB | PB |  | PP | 26.4; 44.7 | 4,0 | 1.3; 9.0 | 100 | 56,0 | 0,0 |
| 69745 | PP | PP | PP |  | PB | PB | PB | PB | PB | 24.6; 25.7 | 11,0 | 5.9; 18.1 | 100 | 64,0 | 0,0 |
| 69744 | PP | PP |  | PB | PB | PB | PB | PB | PB | 18.0; 24.6 | 8,0 | 3.7; 14.4 | 100 | 65,0 | 0,0 |
| 69934 | PB | PB | PB | PB | PB | PB | PB | PB | PB | 0.0; 0.0 | 10,0 | 5.1; 16.9 | 100 | 68,0 | 0,0 |
| 69935 | PB | PB | PB | PB | PB | PB | PB | PB | PB | 0.0; 0.0 | 9,0 | 4.4; 15.6 | 100 | 64,0 | 0,0 |
| 69878 | PB | PB | PB | PB | PB | PB | PB | PB | PB | 0.0; 0.0 | 8,0 | 3.7; 14.4 | 100 | 65,0 | 0,0 |
| 69627 | PB | PB | PB | PB | PB | PB | PB | PB | PB | 0.0; 0.0 | 8,0 | 3.7; 14.4 | 100 | 64,0 | 0,0 |
| 69467 | PB | PB | PB | PB | PB | PB | PB | PB | PB | 0.0; 0.0 | 7,0 | 3.1; 13.1 | 100 | 58,0 | 0,0 |
| 69626 | PB | PB | PB | PB | PB | PB | PB | PB | PB | 0.0; 0.0 | 3,0 | 0.8; 7.6 | 100 | 63,0 | 0,0 |
| 69780 | PB | PB | PB | PB | PB | PB | PB | PB | PB | 0.0; 0.0 | 1,0 | 0.1; 4.3 | 100 | 60,0 | 0,0 |
| 69572 | PB | PB | PB | PB | PB | PB | PB |  | PP | 0.7; 14.4 | 8,0 | 3.7; 14.4 | 100 | 57,0 | 0,0 |
| 37007 | PB | PB | PB | PB | PB | PB | PB |  | PP | 0.7; 14.4 | 4,0 | 1.2; 9.1 | 98 | 69,0 | 0,0 |
| 69625 | PB | PB | PB | PB | PB | PB | PB |  | PP | 0.7; 14.4 | 1,0 | 0.1; 4.3 | 100 | 65,0 | 0,0 |
| 69877 | PB | PB | PB | PB | PB | PB |  | PP | PP | 14.4; 25.7 | 1,0 | 0.1; 4.3 | 100 | 56,0 | 0,0 |
| 69781 | PB | PB | PB |  | PP | PP | PP | PP |  | 100.7; 119.0 | 3,0 | 0.8; 7.6 | 100 | 62,0 | 0,0 |

The minimal and maximal length of PWD/PWD interval was determined by positions of the DNA markers used for genotyping. White and orange intervals are heterozygous PWD/B6 and homozygous PWD/PWD, respectively. Recombination breaks are located in gray intervals.
Sperm count refers to the number of sperm cells in both epididymis, TW is the wet weight of paired testes in milligrams.

**Table supplement 6.** The effect of the size and location of PWD/PWD consubspecific intervals on asynapsis of Chr 15 and on fertility parameter

| Male ID | Distance from centromere of Chr 15 (total length 104.0 Mb) |  |  |  |  |  |  |  |  |  | PWD/PWD length | Asynapsis of Chr 15 |  | Cells analyzed | Testes weight | Sperm count |
| --- | --- | --- | --- | --- | --- | --- | --- | --- | --- | --- | --- | --- | --- | --- | --- | --- |
|  | 9,3 | 40,0 | 63,0 | 70,2 | 97,0 | 97,8 | 100,6 | 101,9 | 102,8 | 103,4 | Mb, range | % | 95% CI | N | mg | 10<sup>6</sup> |
| 69571 | PP | PP |  | PB | PB | PB | PB | PB | PB | PB | 40.0; 63.0 | 2,0 | 0.3; 6.0 | 100 | 69,0 | 0,0 |
| 69623 | PP |  | PB | PB | PB | PB | PB | PB | PB | PB | 9.3; 40.0 | 2,0 | 0.3; 6.0 | 100 | 65,0 | 0,0 |
| 69783 | PB | PB | PB | PB | PB | PB | PB | PB | PB | PB | 0.0; 0.0 | 3,0 | 0.8; 7.6 | 100 | 56,0 | 0,0 |
| 69784 | PB | PB | PB | PB | PB | PB | PB | PB | PB | PB | 0.0; 0.0 | 11,0 | 5.9; 18.1 | 100 | 60,0 | 0,0 |
| 69566 | PB | PB | PB | PB | PB | PB | PB | PB | PB | PB | 0.0; 0.0 | 13,0 | 7.4; 20.5 | 100 | 72,0 | 0,0 |
| 69469 | PB | PB | PB | PB | PB | PB | PB | PB | PB | PB | 0.0; 0.0 | 16,0 | 9.7; 24.0 | 100 | 56,0 | 0,0 |
| 69565 | PB | PB | PB | PB | PB | PB | PB | PB | PB | PB | 0.0; 0.0 | 16,0 | 9.7; 24.0 | 100 | 68,0 | 0,0 |
| 69625 | PB | PB | PB | PB | PB | PB | PB | PB | PB | PB | 0.0; 0.0 | 16,0 | 9.7; 24.0 | 100 | 65,0 | 0,0 |
| 69624 | PB | PB | PB | PB | PB | PB | PB | PB | PB | PB | 0.0; 0.0 | 19,0 | 12.1; 27.4 | 100 | 59,0 | 0,0 |
| 69746 | PB | PB | PB | PB | PB | PB | PB | PB | PB | PB | 0.0; 0.0 | 21,0 | 13.8; 29.7 | 100 | 62,0 | 0,0 |
| 69743 | PB | PB | PB | PB | PB | PB | PB | PB | PB | PB | 0.0; 0.0 | 21,0 | 13.8; 29.7 | 100 | 65,0 | 0,0 |
| 69781 | PB | PB | PB | PB | PB | PB | PB |  | PP | PP | 1.2; 2.1 | 24,0 | 16.4; 33.0 | 100 | 62,0 | 0,0 |
| 69878 | PB | PB | PB | PB | PB | PB | PB |  | PP | PP | 1.2; 2.1 | 23,0 | 15.5; 31.9 | 100 | 65,0 | 0,0 |
| 69782 | PB | PB | PB | PB | PB | PB | PB |  | PP | PP | 1.2; 2.1 | 16,0 | 9.7; 24.0 | 100 | 58,0 | 0,0 |
| 69879 | PB | PB | PB | PB |  | PP | PP | PP | PP | PP | 6.2; 7.0 | 0,0 | 0.0; 2.0 | 100 | 58,0 | 0,0 |

The minimal and maximal length of PWD/PWD interval was determined by positions of the DNA markers used for genotyping. White and orange intervals are heterozygous PWD/B6 and homozygous PWD/PWD, respectively. Recombination breaks are located in gray intervals.
Sperm count refers to the number of sperm cells in both epididymis, TW is the wet weight of paired testes in milligrams.

**Table supplement 7.** The effect of the size and location of PWD/PWD consubspecific intervals on asynapsis of Chr 17 and on fertility parameters

| Male ID | Distance from centromere of Chr 17 (total length 95.0 Mb) |  |  |  |  |  |  |  |  |  | PWD/PWD length | Asynapsis of Chr 17 |  | Cells analyzed | Testes weight | Sperm count |
| --- | --- | --- | --- | --- | --- | --- | --- | --- | --- | --- | --- | --- | --- | --- | --- | --- |
|  | 4,8 | 15 | 15,6 | 30,9 | 34,7 | 45,4 | 52,7 | 65,3 | 73,8 | 93,2 | Mb, range | % | 95% CI | N | mg | 10<sup>6</sup> |
| 64979 | PB | PB | PB | PB | PB | PB | PB | PB | PB | PB | 0.0; 0.0 | 18,8 | 12.0; 27.1 | 101 | 72,0 | 0,0 |
| 66707 | PB | PB | PB | PB | PB | PB | PB | PB | PB | PB | 0.0; 0.0 | 14,9 | 8.9; 22.7 | 101 | 63,0 | 0,0 |
| 64855 | PB | PB | PB | PB | PB | PB | PB | PB | PB | PB | 0.0; 0.0 | 14,9 | 8.9; 22.7 | 101 | 73,0 | 0,0 |
| 66954 | PB | PB | PB | PB | PB | PB | PB | PB | PB | PB | 0.0; 0.0 | 14,6 | 8.7; 22.3 | 103 | 71,0 | 0,0 |
| 64574 | PB | PB | PB | PB | PB | PB | PB | PB | PB | PB | 0.0; 0.0 | 11,7 | 6.5; 18.8 | 103 | 68,0 | 0,0 |
| 64977 | PB | PB | PB | PB | PB | PB | PB | PB | PB | PB | 0.0; 0.0 | 11,2 | 6.3; 17.8 | 116 | 63,0 | 0,0 |
| 64718 | PB | PB | PB | PB | PB | PB | PB | PB | PB | PB | 0.0; 0.0 | 12,5 | 6.9; 20.1 | 96 | 68,0 | 0,0 |
| 64849 | PB | PB | PB | PB | PB | PB | PB | PB | PB | PB | 0.0; 0.0 | 7,0 | 3.1; 13.0 | 102 | 76,0 | 0,0 |
| 66515 | PB | PB | PB | PB | PB | PB | PB | PB |  | PP | 1.8; 21.2 | 16,0 | 9.5; 24.4 | 92 | 75,0 | 0,0 |
| 66514 | PB | PB | PB | PB | PB | PB | PB | PB |  | PP | 1.8; 21.2 | 14,0 | 8.1; 21.7 | 100 | 71,0 | 0,0 |
| 64851 | PB | PB | PB | PB | PB | PB | PB | PB |  | PP | 1.8; 21.2 | 10,5 | 5.6; 17.3 | 105 | 69,0 | 0,0 |
| 64719 | PB | PB | PB | PB | PB | PB | PB | PB |  | PP | 1.8; 21.2 | 5,8 | 2.4; 11.3 | 106 | 62,0 | 0,0 |
| 66819 | PB | PB | PB | PB | PB | PB | PB | PB |  | PP | 1.8; 21.2 | 3,4 | 0.9; 8.2 | 100 | 72,0 | 0,0 |
| 66813 | PB | PB | PB | PB | PB | PB | PB |  | PP | PP | 21.2; 29.7 | 1,0 | 0.1; 4.2 | 104 | 68,0 | 0,0 |
| 66876 | PB | PB | PB | PB | PB |  |  | PP | PP | PP | 29.7; 42.3 | 0,0 | 0.0; 2.0 | 97 | 58,0 | 0,0 |
| 64575 | PB | PB | PB | PB | PB |  | PP | PP | PP | PP | 42.3; 49.6 | 0,0 | 0.0; 1.9 | 103 | 73,0 | 0,0 |
| 64922 | PB | PB | PB | PB | PB |  | PP | PP | PP | PP | 42.3; 49.6 | 2,8 | 0.7; 7.1 | 108 | 69,0 | 0,0 |
| 64975 | PP |  | PB | PB | PB | PB | PB | PB | PB | PB | 4.8; 15.0 | 7,9 | 3.7; 14.2 | 101 | 80,0 | 0,0 |
| 66532 | PP |  | PB | PB | PB | PB | PB | PB | PB | PB | 4.8; 15.0 | 6,0 | 2.4; 11.8 | 100 | 72,0 | 0,0 |

The minimal and maximal length of PWD/PWD interval was determined by positions of the DNA markers used for genotyping. White and orange intervals are heterozygous PWD/B6 and homozygous PWD/PWD, respectively. Recombination breaks are located in gray intervals.

**Table supplement 8.** The effect of the size and location of PWD/PWD consubspecific intervals on asynapsis of Chr 18 and on fertility parameters

| Male ID | Distance from centromere of Chr 18 (90.7 Mb) |  |  |  |  | PWD/PWD<br>length | Asynapsis of Chr 18 |  | Cells<br>analyzed | Testes<br>weight | Sperm<br>count |
| --- | --- | --- | --- | --- | --- | --- | --- | --- | --- | --- | --- |
|  | 3,8 | 25,0 | 57,4 | 76,9 | 89,6 | Mb, range | % | 95% CI | N | mg | 10<sup>6</sup> |
| 64574 | PB | PB | PB | PB | PB | 0.0; 0.0 | 30,8 | 22.5; 40.1 | 104 | 68,0 | 0,0 |
| 64851 | PB | PB | PB | PB | PB | 0.0; 0.0 | 21,4 | 14.2; 30.0 | 103 | 69,0 | 0,0 |
| 64719 | PB | PB | PB | PB | PB | 0.0; 0.0 | 19,3 | 12.7; 27.4 | 109 | 62,0 | 0,0 |
| 64976 | PB | PB | PB |  | PP | 1.1; 13.8 | 28,0 | 19.8; 37.3 | 100 | 71,0 | 0,0 |
| 66876 | PB | PB |  | PP | PP | 13.8; 33.3 | 9,4 | 3.9; 18.1 | 64 | 58,0 | 0,0 |
| 66877 | PB | PB |  | PP | PP | 13.8; 33.3 | 1,0 | 0.1; 4.3 | 101 | 64,0 | 0,0 |
| 66532 | PB | PB |  | PP | PP | 13.8; 33.3 | 0,0 | 0.0; 2.5 | 78 | 72,0 | 0,0 |
| 66954 | PB |  | PP | PP | PP | 33.3; 65.7 | 0,0 | 0.0; 1.9 | 102 | 71,0 | 0,0 |
| 66813 | PB |  | PP | PP | PP | 33.3; 65.7 | 0,0 | 0.0; 1.7 | 113 | 68,0 | 0,0 |
| 64922 | PP | PP | PP | PP | PP | 90.7; 90.7 | 1,8 | 0.3; 5.5 | 109 | 69,0 | 0,0 |
| 64718 | PP | PP | PP | PP | PP | 90.7; 90.7 | 0,0 | 0.0; 2.0 | 100 | 68,0 | 0,0 |
| 64975 | PP | PP | PP | PP | PP | 90.7; 90.7 | 0,0 | 0.0; 2.0 | 100 | 80,0 | 0,0 |
| 64855 | PP | PP | PP | PP |  | 76.9; 89.6 | 0,0 | 0.0; 1.9 | 103 | 73,0 | 0,0 |
| 64849 | PP | PP | PP | PP |  | 76.9; 89.6 | 0,0 | 0.0; 2.0 | 100 | 76,0 | 0,0 |
| 64575 | PP | PP | PP | PP |  | 76.9; 89.6 | 2,0 | 0.3; 6.0 | 101 | 73,0 | 0,0 |
| 64968 | PP | PP | PP | PP |  | 76.9; 89.6 | 3,0 | 0.8; 7.6 | 101 | 73,0 | 0,0 |
| 64979 | PP | PP | PP |  | PB | 57.4; 76.9 | 1,0 | 0.1; 4.3 | 100 | 72,0 | 0,0 |
| 66872 | PP | PP | PP |  | PB | 57.4; 76.9 | 2,0 | 0.3; 6.0 | 103 | 73,0 | 0,0 |
| 66514 | PP | PP | PP |  | PB | 57.4; 76.9 | 3,0 | 0.8; 7.6 | 100 | 71,0 | 0,0 |
| 64977 | PP | PP |  | PB | PB | 25.0; 57.4 | 4,8 | 1.7; 10.0 | 104 | 63,0 | 0,0 |
| 66707 | PP |  | PB | PB | PB | 3.8; 25.0 | 4,9 | 1.8; 10.2 | 103 | NA | 0,0 |
| 66815 | PP |  | PB | PB | PB | 3.8; 25.0 | 11,0 | 6.0; 17.8 | 109 | 75,0 | 0,0 |

The minimal and maximal length of PWD/PWD interval was determined by positions of the DNA markers used for genotyping. White and orange intervals are heterozygous PWD/B6 and homozygous PWD/PWD, respectively. Recombination breaks are located in gray intervals.
Sperm count refers to the number of sperm cells in both epididymis, TW is the wet weight of paired testes in milligrams.

**Table supplement 9.**
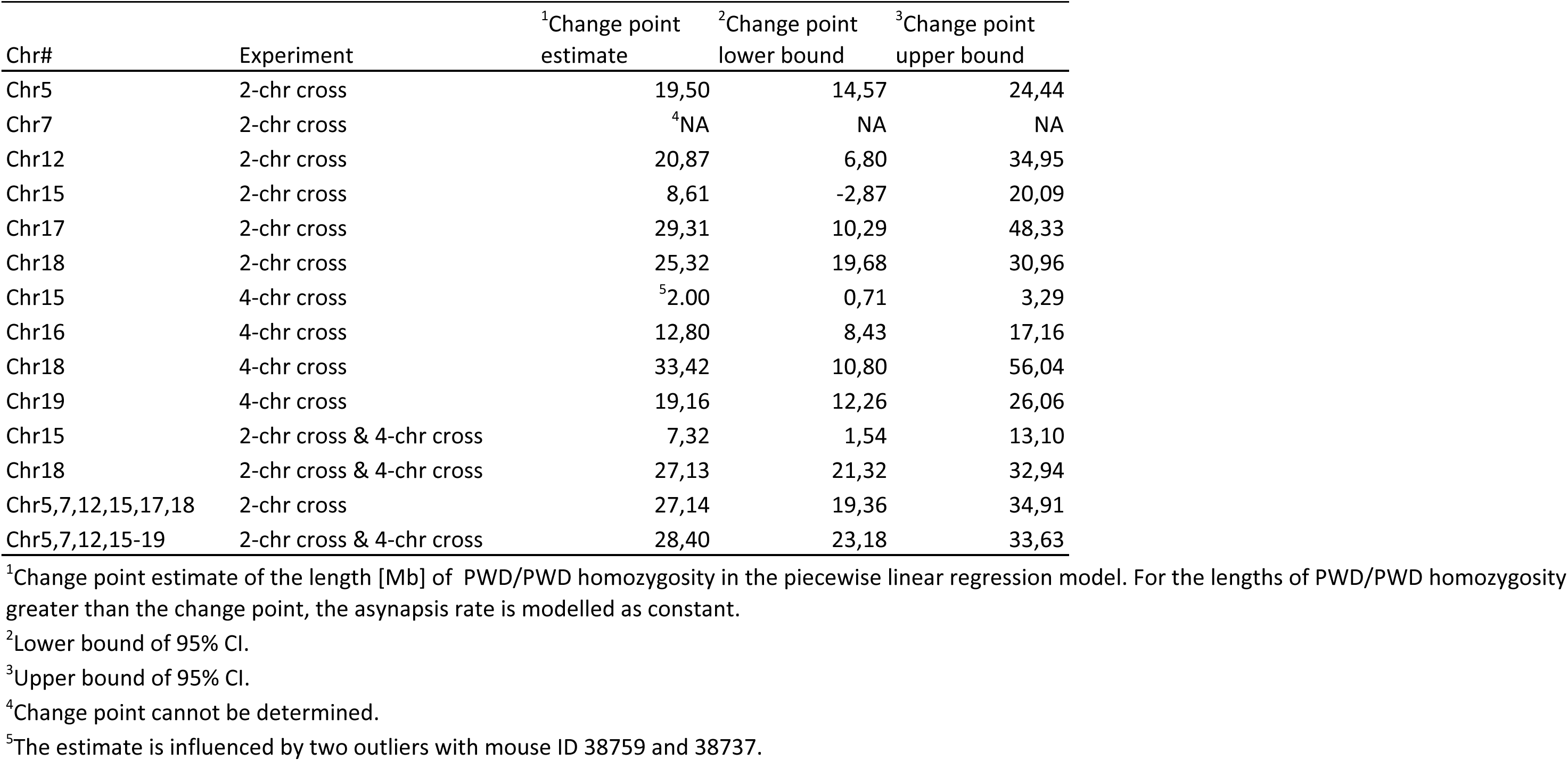
Change point estimates of the minimal length [Mb] of PWD/PWD homozygosity showing detectable affect on synapsis rate

**Table supplement 10.**
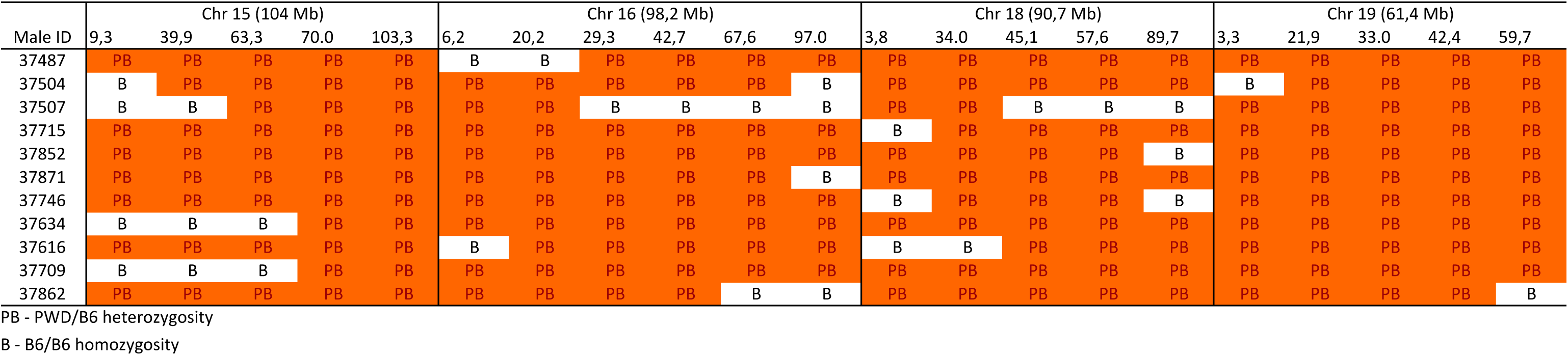
Eleven G3 male parents selected for the 4-chr cross experiment.

**Table supplement 11.** The fertility parameters of hybrids of the 4-chr cross experiment

| Male ID | TW (mg) | Sperm x 10<sup>-6</sup> |
| --- | --- | --- |
| 38754 | 51 | 0,00 |
| 38428 | 57 | 0,00 |
| 38735 | 58 | 0,00 |
| 38088 | 59 | 0,00 |
| 38089 | 60 | 0,00 |
| 38753 | 60 | 0,00 |
| 38726 | 61 | 0,00 |
| 38751 | 61 | 0,00 |
| 38755 | 61 | 0,00 |
| 38435 | 62 | 0,00 |
| 38090 | 63 | 0,00 |
| 38085 | 64 | 0,00 |
| 38087 | 64 | 0,00 |
| 38448 | 64 | 0,00 |
| 38738 | 65 | 0,00 |
| 38739 | 65 | 0,00 |
| 38711 | 65 | 0,00 |
| 38728 | 65 | 0,00 |
| 38822 | 65 | 0,00 |
| 38430 | 66 | 0,00 |
| 38432 | 66 | 0,00 |
| 38710 | 66 | 0,00 |
| 38084 | 67 | 0,00 |
| 38434 | 67 | 0,00 |
| 38814 | 67 | 0,00 |
| 38752 | 68 | 0,00 |
| 38783 | 68 | 0,00 |
| 38817 | 68 | 0,00 |
| 38758 | 69 | 0,00 |
| 38821 | 69 | 0,00 |
| 38740 | 70 | 0,00 |
| 38766 | 70 | 0,00 |
| 38813 | 70 | 0,00 |
| 38426 | 71 | 0,00 |
| 38468 | 71 | 0,00 |
| 38765 | 72 | 0,00 |
| 38767 | 73 | 0,00 |
| 38819 | 73 | 0,00 |
| 38792 | 74 | 0,00 |
| 38818 | 74 | 0,00 |
| 38820 | 74 | 0,00 |
| 38815 | 75 | 0,00 |
| 75890 | 68 | 0,00 |
| 79414 | 65 | 0,00 |
| 38712 | 76 | 0,00 |
| 38733 | 73 | 0,01 |
| 38732 | 63 | 0,01 |
| 38561 | 78 | 0,01 |
| 38713 | 77 | 0,03 |
| 38727 | 62 | 0,03 |
| 38757 | 72 | 0,04 |
| 38427 | 64 | 0,06 |
| 38429 | 64 | 0,09 |
| 38425 | 73 | 0,10 |
| 38566 | 63 | 0,17 |
| 38734 | 66 | 0,21 |
| 38086 | 73 | 0,23 |
| 38725 | 69 | 0,29 |
| 38852 | 75 | 0,34 |
| 38842 | 89 | 0,43 |
| 38759 | 81 | 0,60 |
| 38858 | 73 | 0,74 |
| 38763 | 81 | 1,02 |
| 38736 | 81 | 1,10 |
| 38764 | 84 | 1,10 |
| 38741 | 81 | 1,20 |
| 38801 | 83 | 1,51 |
| 38565 | 70 | 1,61 |
| 38794 | 90 | 1,80 |
| 38737 | 94 | 1,90 |
| 75889 | 90 | 2,10 |
| 38692 | 89 | 2,20 |
| 79415 | 83 | 2,20 |
| 38724 | 71 | 2,80 |
| 79417 | 90 | 2,80 |
| 79412 | 103 | 3,90 |
| 38433 | 78 | 4,20 |
| 38816 | 106 | 4,20 |
| 38756 | 85 | 4,70 |
| 38650 | 82 | 5,40 |
| 38652 | 90 | 5,60 |
| 38431 | 81 | 6,40 |
| 79416 | 95 | 6,40 |
| 38555 | 98 | 6,87 |
| 38793 | 117 | 10,20 |
| 75891 | 131 | 12,40 |
| 79413 | 108 | 13,70 |

<sup>†</sup>The minimal and maximal length of PWD/PWD interval was determined by positions of the DNA markers used for genotyping. White and orange intervals are heterozygous PWD/B6 and homozygous PWD/PWD, respectively. Recombination breaks are located in gray intervals.
<sup>2</sup>Sperm count refers to the number of sperm cells in both epididymis.
<sup>3</sup>The average between minimum and maximum PWD/PWD consubspecific sequence is given.

**Table supplement 12.**
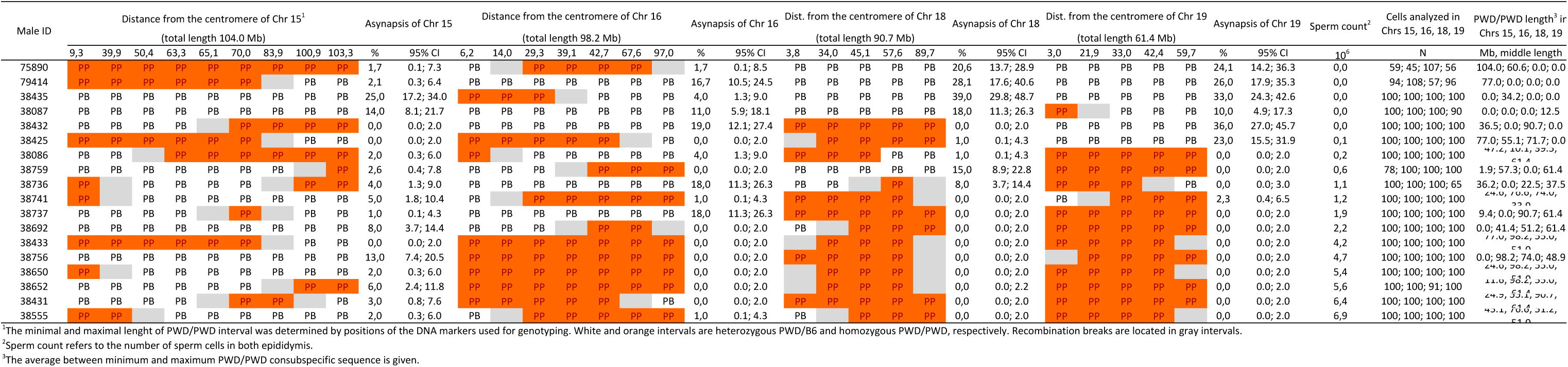
Four-chr cross. The effect of the size and location of PWD/PWD consubspecific intervals on asynapsis of Chrs 15, 16, 18 and 19 and on fertility parameters.

**Table supplement 13.** Four-chr cross experiment. The relation between observed and calculated rate of pachytene asynapsis and fertility parameters.

| Male ID | Testes (mg) | Sperm No<br>$\times 10^{-6}$ | Frequency of Chr synapsed | | | | Probability<br>of all 4 chr<br>synapsed | Probability<br>of all 19<br>synapsed | Probability<br>of asynaptic<br>pachy | Obs ratio of<br>asynaptic<br>pachy | Obs ratio of<br>all 19<br>synapsed |
| --- | --- | --- | --- | --- | --- | --- | --- | --- | --- | --- | --- |
|  |  |  | 15 | 16 | 18 | 19 |  |  |  |  |  |
| 38435 | 62 | 0 | 0,750 | 0,960 | 0,610 | 0,670 | 0,294 | 0,079 | 0,921 | 0,790 | 0,210 |
| 38087 | 64 | 0 | 0,860 | 0,890 | 0,820 | 0,900 | 0,565 | 0,151 | 0,849 | 0,610 | 0,390 |
| 75414 | 65 | 0 | 0,979 | 0,820 | 0,719 | 0,740 | 0,427 | 0,114 | 0,886 | 0,660 | 0,340 |
| 38432 | 66 | 0 | 1,000 | 0,810 | 1,000 | 0,640 | 0,518 | 0,138 | 0,862 | 0,650 | 0,350 |
| 75890 | 68 | 0 | 0,983 | 0,959 | 0,793 | 0,759 | 0,567 | 0,151 | 0,849 | 0,630 | 0,370 |
| 38425 | 73 | 0,1 | 1,000 | 1,000 | 0,990 | 0,770 | 0,762 | 0,204 | 0,796 | 0,580 | 0,420 |
| 38086 | 73 | 0,23 | 0,980 | 0,960 | 0,990 | 1,000 | 0,931 | 0,249 | 0,751 | 0,360 | 0,640 |
| 38759 | 81 | 0,6 | 0,970 | 1,000 | 0,850 | 1,000 | 0,825 | 0,220 | 0,780 | 0,490 | 0,510 |
| 38736 | 81 | 1,1 | 0,960 | 0,820 | 0,920 | 1,000 | 0,724 | 0,193 | 0,807 | 0,480 | 0,520 |
| 38741 | 81 | 1,2 | 0,950 | 0,990 | 1,000 | 0,980 | 0,922 | 0,246 | 0,754 | 0,480 | 0,520 |
| 38737 | 94 | 1,9 | 0,990 | 0,820 | 1,000 | 1,000 | 0,812 | 0,217 | 0,783 | 0,400 | 0,600 |
| 38692 | 89 | 2,2 | 0,920 | 1,000 | 1,000 | 1,000 | 0,920 | 0,246 | 0,754 | 0,280 | 0,720 |
| 38433 | 78 | 4,2 | 1,000 | 1,000 | 1,000 | 1,000 | 1,000 | 0,267 | 0,733 | 0,280 | 0,720 |
| 38756 | 85 | 4,7 | 0,870 | 1,000 | 1,000 | 1,000 | 0,870 | 0,232 | 0,768 | 0,390 | 0,610 |
| 38650 | 82 | 5,4 | 0,980 | 1,000 | 1,000 | 1,000 | 0,980 | 0,262 | 0,738 | 0,360 | 0,640 |
| 38652 | 90 | 5,6 | 0,940 | 1,000 | 1,000 | 1,000 | 0,940 | 0,251 | 0,749 | 0,330 | 0,670 |
| 38431 | 81 | 6,4 | 0,970 | 1,000 | 1,000 | 1,000 | 0,970 | 0,259 | 0,741 | 0,460 | 0,540 |
| 38555 | 98 | 6,9 | 0,980 | 0,960 | 1,000 | 1,000 | 0,941 | 0,251 | 0,749 | 0,255 | 0,745 |

**Table supplement 14.** Selected SSLP markers polymorphic between B6 and PWD.

| Chr No. | SSLP marker | Mbp |
| --- | --- | --- |
| 5 | D5Mit346 | 4,88 |
| 5 | D5Mit49 | 5,20 |
| 5 | D5Mit178 | 9,83 |
| 5 | D5Mit349 | 24,97 |
| 5 | D5Mit386 | 28,10 |
| 5 | D5Mit106 | 44,99 |
| 5 | D5Mit302 | 61,43 |
| 5 | D5Mit303 | 64,10 |
| 5 | D5Mit277 | 107,06 |
| 5 | D5Mit136 | 118,73 |
| 5 | D5Mit138 | 125,10 |
| 5 | D5Mit140 | 126,95 |
| 5 | D5Mit59 | 127,72 |
| 5 | D5Mit63 | 132,13 |
| 5 | D5Mit326 | 137,43 |
| 5 | D5Mit102 | 151,35 |
| 7 | D7Mit21 | 3,27 |
| 7 | D7Mit191 | 18,89 |
| 7 | D7Mit180 | 24,63 |
| 7 | D7Mit265 | 25,68 |
| 7 | D7Mit246 | 30,30 |
| 7 | D7Mit31 | 87,49 |
| 7 | D7Mit66 | 119,89 |
| 7 | D7Mit71 | 131,03 |
| 7 | D7Mit259 | 144,76 |
| 12 | D12Mit37 | 5,37 |
| 12 | D12Mit215 | 7,61 |
| 12 | D12Mit49 | 15,71 |
| 12 | D12Mit11 | 22,39 |
| 12 | D12Mit136 | 30,17 |
| 12 | D12Mit147 | 35,72 |
| 12 | D12Mit285 | 54,65 |
| 12 | D12Mit118 | 91,51 |
| 12 | D12Mit7 | 103,40 |
| 12 | D12Mit20 | 113,36 |
| 15 | D15Mit175 | 9,27 |
| 15 | D15Mit130 | 20,81 |
| 15 | D15Mit9 | 26,04 |
| 15 | D15Mit138 | 40,03 |
| 15 | D15Mit113 | 50,40 |
| 15 | D15Mit270 | 63,52 |
| 15 | D15Mit234 | 65,07 |
| 15 | D15Mit67 | 70,20 |
| 15 | D15Mit215 | 83,89 |
| 15 | D15Mit161 | 97,01 |
| 15 | D15Mit43 | 97,99 |
| 15 | D15Mit97 | 100,80 |
| 15 | D15Mit40 | 102,03 |
| 15 | D15Mit16 | 102,99 |
| 15 | D15Mit79 | 103,47 |
| 16 | D16Mit55 | 6,24 |
| 16 | D16Mit165 | 14,00 |
| 16 | D16Mit29 | 20,20 |
| 16 | D16Mit57 | 29,30 |
| 16 | D16Mit12 | 39,12 |
| 16 | D16Mit83 | 42,81 |
| 16 | D16Mit76 | 67,72 |
| 16 | D16Mit71 | 96,91 |
| 17 | D17Mit19 | 4,81 |
| 17 | D17M334 | 15,00 |
| 17 | D17M634 | 15,60 |
| 17 | D17Mit81 | 30,90 |
| 17 | D17Mit33 | 34,74 |
| 17 | D17Mit50 | 45,36 |
| 17 | D17Mit139 | 52,66 |
| 17 | D17Mit152 | 65,34 |
| 17 | D17Mit93 | 73,80 |
| 17 | D17Mit122 | 83,06 |
| 17 | D17Mit155 | 84,50 |
| 17 | D17Mit123 | 93,20 |
| 18 | D18Mit66 | 3,84 |
| 18 | D18Mit22 | 24,95 |
| 18 | D18Mit84 | 33,83 |
| 18 | D18Mit70 | 34,85 |
| 18 | D18Mit149 | 45,00 |
| 18 | D18Mit53 | 52,83 |
| 18 | D18Mit55 | 53,16 |
| 18 | D18Mit54 | 53,43 |
| 18 | D18Mit124 | 57,46 |
| 18 | D18Mit184 | 66,82 |
| 18 | D18Mit209 | 66,97 |
| 18 | D18Mit185 | 69,03 |
| 18 | D18Mit7 | 76,85 |
| 18 | D18Mit25 | 89,57 |
| 19 | D19Mit32 | 3,28 |
| 19 | D19Mit61 | 15,53 |
| 19 | D19Mit128 | 17,25 |
| 19 | D19Mit96 | 21,84 |
| 19 | D19Mit46 | 32,94 |
| 19 | D19Mit11 | 42,40 |
| 19 | D19Mit71 | 59,60 |

